# Click-chemistry-aided quantitation and sequencing of oxidized guanines and apurinic sites uncovers their transcription-linked strand bias in human cells

**DOI:** 10.1101/2024.07.21.604463

**Authors:** Vakil Takhaveev, Nikolai J.L. Püllen, Navnit K. Singh, Sabrina M. Huber, Stefan Schauer, Hailey L. Gahlon, Anna R. Poetsch, Shana J. Sturla

**Author notes:** These authors contributed equally (Vakil Takhaveev, Nikolai J.L. Püllen).

## Abstract

DNA modifications drive aging, neurodegeneration, carcinogenesis, and chemotherapy drug action. To understand the functional genomic roles of DNA modifications, it is critical to accurately map their diverse chemical forms with single-nucleotide precision in complex genomes, but it remains challenging. Click-code-seq is a click-chemistry-aided single-nucleotide-resolution strategy for guanine-oxidation mapping, used in yeast DNA but having poor applicability to human genomes. Here, we upgraded click-code-seq to enable its first application for sequencing DNA oxidation and depurination in human genomes. For this, we developed a companion fluorescence assay, click-fluoro-quant, to rapidly quantify different common DNA modifications, and devised novel adapters to minimize false modification detection and assess modification frequency in cell populations. We uncovered that endogenous DNA oxidation in a human cell line has a highly similar pattern to cancer mutational signatures associated with reactive oxygen species. We established that the DNA-alkylating chemotherapy drug irofulven preferentially induces depurination in ApA dimers and promoter regions. Intriguingly, we revealed that oxidized guanines and apurinic sites, both irofulven-induced and endogenous, are depleted in gene transcribed strands, and the strand bias widens with increasing gene expression. This work substantially advances click-code-seq for deciphering the impacts of key modifications in human DNA on cellular physiology and toxicological responses.

## Introduction

Genomic DNA can undergo chemically diverse modifications due to various endogenous factors such as replication errors, reactive oxygen species (ROS) and spontaneous hydrolysis, and due to reactions with exogenous agents like environmental alkylating chemicals, drugs, and UV and ionizing radiation^1–4^. Apart from enzymatically imprinted epigenetic modifications, the most abundant endogenous forms include DNA breaks, abasic sites (AP sites), 8-oxoguanines (8-oxoG), lipid peroxidation products and alkylation adducts, which occur at a frequency of about 10^5^ events per mammalian cell per day^3–5^. DNA modifications are causally linked to most phenotypic aspects of aging through the activation of DNA damage signaling^6,7^ and are implicated in neurodegenerative deseases^8,9^. Moreover, they drive carcinogenesis as precursors for mutations^10–13^. Conversely, many anticancer drugs modify DNA to target rapidly dividing cancer cells by inhibiting DNA replication and transcription^14,15^. Beyond cytosine methylation and its derivatives, DNA modifications such as breaks, AP sites and 8-oxoG are also emerging as putative new types of epigenetic marks, with their association to promoters appearing to control gene expression^16–20^. While DNA breaks have been mapped with single-nucleotide resolution in mammalian genomes via a range of experimental methods^15,21–26^, there are, however, limited data concerning the precise locations of AP sites and 8-oxoG in large genomes.

AP sites have been mapped at single-nucleotide resolution in human cells in two studies^27,28^, with one of them locating these DNA modifications also in mouse tissues^28^. AP sites were strongly enriched in purines, however, guanine appeared to be the most frequently depurinated nucleotide in one work^27^, whereas it was adenine in the other^28^. Additionally, while, according to the first work, AP sites were stochastically distributed across the genome without site specificity at the single-nucleotide level^27^, it was found in the second study that AP sites accumulated at single-nucleotide hotspots shared across samples, were associated with sequence variants and, in mouse tissues, occurred more frequently in highly expressed genes, with a bias towards the non-template strand^28^. The discrepancies regarding the precise location of AP sites between these studies may be due to differences in the methods used: snAP-seq, which relies on chemical tagging of the reactive aldehyde moiety at AP sites^27^, and SSiNGLe-AP, which involves endonuclease-mediated conversion of AP sites to 3’-OH termini followed by capture through polyA tailing^28^.

To the best of our knowledge, only one study has mapped 8-oxoG at single-nucleotide resolution in a large genome, namely in a human cancer cell line with and without exposure to the oxidant potassium bromate^29^. Therein 8-oxoG was observed to be depleted in GC-rich regions and promoters, partly due to reduced 8-oxoG occurrence at G-quadruplexes^29^. The method used, CLAPS-seq^29^, combined chemical labeling of 8-oxoG from the 150-base-pair-resolution OG-seq assay^30^ with a DNA polymerase stalling strategy to mark modification locations at single-nucleotide resolution^31–33^. The other available single-nucleotide-precision method, click-code-seq, has only been used to map endogenous guanine oxidation in yeast^34^, which has a genome over 256 times shorter than the human genome (12.1 million vs. 3.1 billion base pairs). Consistent with the CLAPS-seq^29^ data and measurements in an X-ray-exposed human cancer cell line using the 250-base-pair-resolution method AP-seq^35^, oxidized guanines were depleted in gene promotor regions of the yeast genome^34^. In click-code-seq, a guanine-oxidation site is converted to a one-nucleotide gap with base excision repair proteins, and the gap is filled with a synthetic *O*-3′-propargyl modified nucleotide. This 3′-alkynyl DNA is then tagged by the 5′-azido-modified code sequence via a copper(I)-catalyzed click reaction, generating a biocompatible triazole-linked DNA^36^ and allowing single-nucleotide-level detection of the original guanine-oxidation site in DNA sequencing^34^. The click partners can also be reversed using commercially available 3′-azido-2′,3′-dideoxynucleotides and a 5′-alkynyl-modified code sequence, facilitating broader applications that avoid custom chemical synthesis^37^. The click-code-seq-derived map of oxidized guanines in the yeast genome, however, contradicted promoter enrichment in 8-oxoG as observed in a mouse cell line^30^ and mouse tissues^38^ by OG-seq, and in human non-cancerous and mouse cell lines by OxiDIP-Seq^39^. Nonetheless, both OG-seq and OxiDIP-Seq have a resolution of over one hundred base pairs^30,38,39^. Thus, the limited single-nucleotide-precision data on AP sites and guanine oxidation are contradictory among themselves as well as when compared to previous lower-resolution measurements.

Significant challenges remain in high-precision genomic mapping of AP sites and 8-oxoG. First, naturally occurring DNA modifications can mask the locations of related target forms. To address this, in snAP-seq, aldehyde moieties of AP sites were selectively chemically tagged over 5-formylcytosine and 5-formyluracil, which can occur more frequently than AP sites^27^. On the other hand, in the AP-site mapping by SSiNGLe-AP, pre-existing 3’-OH breaks were blocked by incorporating biotinylated ddCTP, allowing for the necessary endonuclease-based recognition of AP sites via their conversion to 3’-OH breaks for further tagging^28^. In CLAPS-seq^29^, although the chemical labelling that involves oxidation by K2IrBr6 and addition to an amine-conjugated biotin is highly specific for 8-oxoG^30^, the DNA polymerase stalling strategy relied upon to mark the exact 8-oxoG sites may be confounded by stalls caused by AP sites present in enriched DNA fragments. A second issue is that DNA breaks, AP sites, and oxidized bases can arise as artifacts of sample preparation, caused by pipette shearing, sonication, spontaneous depurination or DNA oxidation^40–44^. While in CLAPS-seq 8-oxoG labelling is performed immediately after DNA extraction to minimize the mapping of artefactual oxidation^29^, snAP-seq includes sonication prior to chemical labelling^27^, and SSiNGLe-AP involves high temperatures before affinity capture^28^, making the methods susceptible to artefactual depurination. A third issue is the potential for spurious priming during DNA-library amplifications, limiting the accurate identification of low abundance modifications especially in large genomes. Finally, current approaches cannot distinguish whether a DNA modification of interest is present in one or multiple cells within a population at a specific genomic site. These problems may contribute to inaccurate results of DNA-modification sequencing and explain inconsistent findings across studies involving different methods.

Here, we comprehensively upgraded the click-chemistry-aided DNA-modification-sequencing method click-code-seq, previously introduced to map oxidation in yeast genome^34^ and bacterial plasmid^37^ DNA, to accurately locate oxidized guanines and AP sites in the human genome at single-nucleotide resolution. First, we developed a coupled fluorescence-based assay termed click-fluoro-quant (Fig. 1a) to rapidly (∼3.5 h) quantify DNA breaks, AP sites or guanine oxidation occurring endogenously or induced by exposure to potassium bromate, UVA irradiation or the chemotherapeutic drug^45,46^ irofulven in human cancer and non-cancerous cell lines. Using click-fluoro-quant in turn to optimize click-code-seq, we found that blocking background modification sites is more efficient than repairing them and established a suitably late stage in the library-preparation process to use sonication while preventing tagging of artefactual modifications. Second, we designed a new type of sequencing adapter – called Modified DNA Identifier Sequence, or MoDIS, – to ligate target sites, distinguishing them from library-amplification artefacts and assessing site-specific DNA-modification frequencies in a population of cells. Employing these new strategies to increase the specificity of click-code-seq, we determined the trinucleotide patterns of endogenous DNA oxidation in a human cancer cell line and compared them with known mutational signatures found in human cancers. We next characterized the genome-wide landscapes of AP sites occurring endogenously or induced by irofulven in a human cancer cell line. Further, we integrated the generated DNA-oxidation and AP-site maps with gene-expression data, uncovering evidence for transcription-coupled DNA repair. Finally, we revealed a strand-specific asymmetry of modifications in the mitochondrial DNA (mtDNA).

**Fig. 1.**
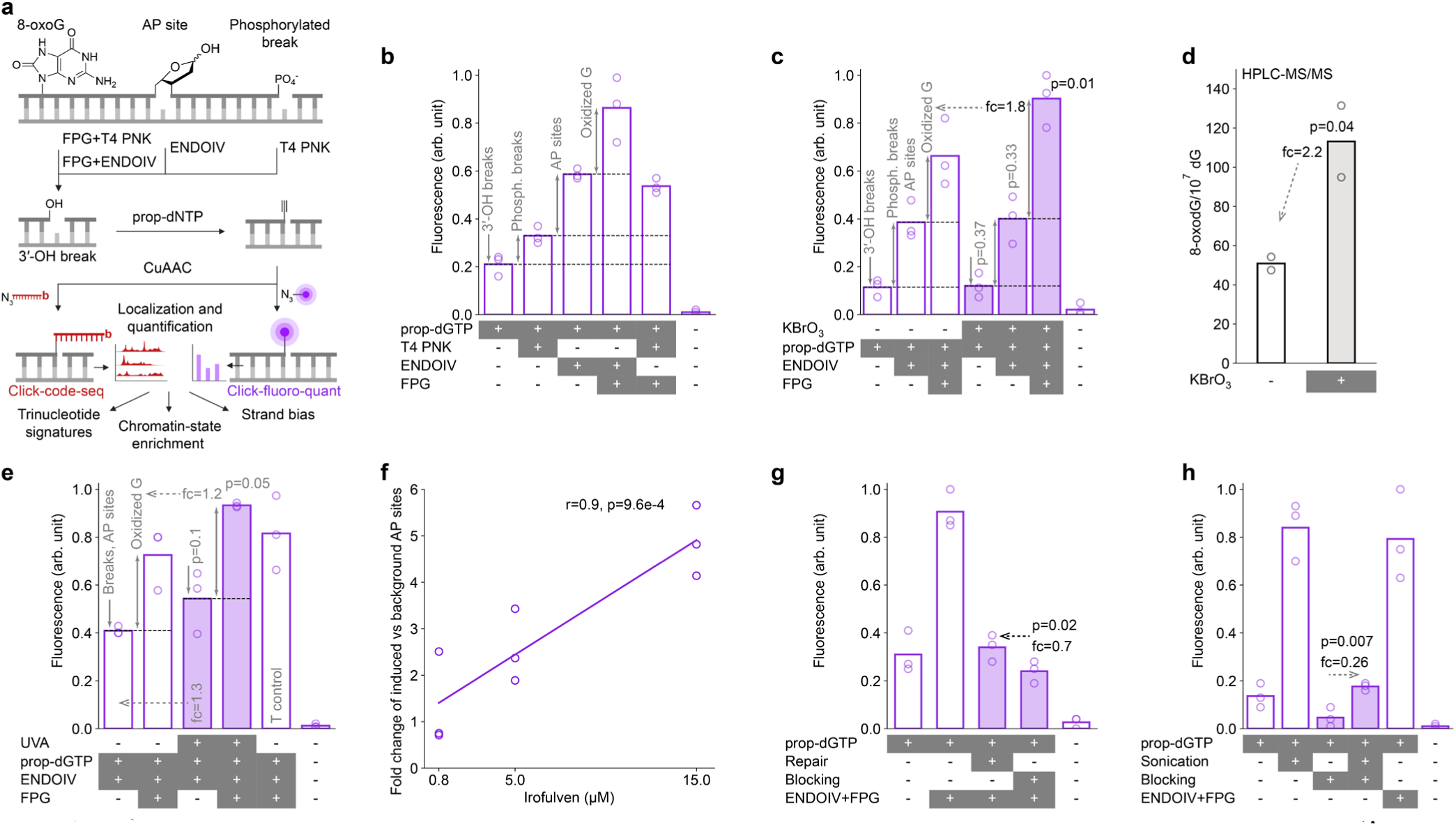
Click-fluoro-quant can be applied to quantify DNA oxidation, AP sites and breaks in genomic DNA. **a** Click-code-seq: clicking DNA-modification sites with a code sequence followed by sequencing; click-fluoro-quant: clicking DNA-modification sites with a fluorophore followed by rapid fluorescence-based quantitation. 8-oxoguanine (8-oxoG), abasic (AP) sites or phosphorylated breaks are converted to 3′-OH-ended breaks via the indicated enzymes (FPG, formamidopyrimidine DNA glycosylase; ENDOIV, endonuclease IV; T4 PNK, T4 polynucleotide kinase), or pre-existing 3′-OH-ended breaks are used. For subsequent ligation, alkyne-modified nucleotides 3’-(*O*-propargyl)-dNTPs (prop-dNTPs) are added to these 3′-OH sites. The alkyne is conjugated via copper click chemistry (CuAAC) with an azide-fluorophore in click-fluoro-quant or a sequencing adapter called Modified DNA Identifier Sequence (MoDIS, red oligonucleotide with b, a biotin group) in click-code-seq. Three directions of genome-wide DNA-modification analysis performed in this work are indicated. **b** Endogenous DNA modifications in HAP1-cell gDNA measured using click-fluoro-quant. **c** Click-fluoro-quant assessment of DNA modifications in HAP1 cells exposed to 50 mM KBrO3 for 30 min. **d** HPLC-MS/MS quantification of 8-oxoG in gDNA of HAP1 cells exposed to 50 mM KBrO3 for 30 min. **e** Click-fluoro-quant assessment of DNA modifications in BJ-5ta cells exposed to 10 J/cm^2^ UVA. T control: cells always stayed in T=37 °C incubator. **f** Click-fluoro-quant assessment of induced AP sites in U2OS cells after 4 h exposure to irofulven. Markers: biological replicates (n=3), each of which included three drug concentrations and a vehicle control by which the plotted values are normalized. r and p: Pearson’s correlation coefficient with respective p-value. **g** Click-fluoro-quant assessment of repair and blocking strategies to remove background DNA modifications in HAP1 gDNA. Repair: nucleotide addition and ligation. Blocking: ddNTP insertion. Before both repair and blocking, 3’-OH groups were released from AP sites and phosphorylated breaks using ENDOIV. **h** Click-fluoro-quant assessment of sonication-induced artifactual DNA modifications with and without ddNTP blocking in HAP1 gDNA. In panels **b-e** and **g-h**, bar: mean of n=3 (click-fluoro-quant) or n=2 (HPLC-MS/MS) biological replicates (markers); p: p-value of the one-tailed (in **c-e**, HA: greater; in **g-h**, HA: less) paired t-test between the exposed and unexposed samples treated with the same set of enzymes (**c-e**) or between the arrow-connected samples (**g-h**); fc: fold change of a specific group of DNA modifications relative to the indicated condition.

## Results

### Rapid quantitation of guanine oxidation, AP sites and breaks in oligonucleotides and genomic DNA by click-fluoro-quant

To conveniently optimize library preparation strategies for DNA-modification-sequencing methods, in particular to remove non-target background and artefactual modifications, we developed a simple fluorescence-based modification-tagging assay, click-fluoro-quant, that we used to compare quantities of 8-oxoG, AP sites and breaks in libraries pre-treated with various protocols. To first establish the method, using a double-stranded 30-mer DNA oligonucleotide with 8-oxoG at position 15, we converted the 8-oxoG to a dephosphorylated 3′-OH site with formamidopyrimidine DNA glycosylase (FPG) and T4 polynucleotide kinase (T4 PNK). We filled this site with 3′-(*O*-propargyl)-dGTP (prop-dGTP) using Therminator IX^47^ and eventually labeled it with AF594-picolyl-azide via copper(I)-catalyzed azide-alkyne cycloaddition (CuAAC, Fig. 1a). The resulting AF594-labeled 15-mer fragment was confirmed by polyacrylamide gel electrophoresis (Supplementary Fig. 1a), and we established a linear correlation between the input concentration of the 8-oxoG-containing oligonucleotide and fluorescence intensity (Supplementary Fig. 1b). We used the same approach, but with prop-dATP, to effectively label AP-site-mimicking tetrahydrofuran in an oligonucleotide by treating it with endonuclease IV (ENDOIV) (Supplementary Fig. 1c). As a result, we could essentially replace sequence bar coding with fluorescence tagging as a confirmation prior to implementing sequencing.

We used the click-fluoro-quant method to distinguish and measure oxidized bases, AP sites and breaks in human genomic DNA (gDNA), by employing corresponding enzymes to covert each modification selectively into 3′-OH groups for subsequent fluorescent labeling (Fig. 1a). When gDNA from the human chronic myeloid leukemia cell line HAP1 was treated with T4 PNK to cleave 3′-phosphorylated DNA breaks, we observed a 1.6-fold increase in fluorescence compared to untreated gDNA where only pre-existing 3′-OH-ended DNA breaks and termini were labeled (Fig. 1b). When the gDNA was treated with ENDOIV, which has both AP-lyase and phosphatase activities^48^, we observed a 2.8-fold increase in fluorescence compared to untreated gDNA and a 1.8-fold increase compared to the T4 PNK-treated samples. The difference between the fluorescence increases for ENDOIV vs. T4 PNK treatments reflects AP sites, accounting for 44% of the signal in ENDOIV-treated gDNA (Fig. 1b). When gDNA was treated with combination of FPG and ENDOIV, which together recognize oxidized bases, AP sites and phosphorylated breaks, we observed an over-4-fold increase in fluorescence compared to when there was no enzymatic treatment (Fig. 1b). This fluorescence signal was 1.6-fold higher than when gDNA was treated with combination of FPG and T4 PNK (Fig. 1b), suggesting that ENDOIV more efficiently cleaved the AP sites produced by FPG.

Having established a selective process for measuring oxidized bases, AP sites or breaks, we exposed cells to representative stressors to assess the potential of click-fluoro-quant for revealing their effects. Thus, HAP1 cells were exposed to the oxidant potassium bromate (KBrO3)^49^. There was no observed change in the levels of breaks or AP sites (Fig. 1c, no enzyme or ENDOIV treatment), however, we observed a 1.8-fold increase of oxidized sites (Fig. 1c, signal differential between ENDOIV+FPG vs. ENDOIV treatments), consistent with a previous report^49^ and quantification of 8-oxoG under the same conditions by HPLC-MS/MS (2.2-fold increase, Fig. 1d). Next, immortalized human skin BJ-5ta cells were irradiated with a dose of 10 J/cm^2^ UVA, leading to a 1.3-fold increase in breaks plus AP sites (Fig. 1e, ENDOIV), and a 1.2-fold increase in DNA oxidation (Fig. 1e, differential between ENDOIV+FPG vs. ENDOIV). Finally, we used the chemotherapeutic drug irofulven^50,51^ to induce depurination in gDNA of human osteosarcoma-derived U2OS cells and identified a drug-concentration-dependent increase in the abundance of induced AP sites (Fig. 1f) but not breaks (Supplementary Fig. 1d). These data support the versatility of using click-fluoro-quant to determine alterations in the targeted modifications induced by chemical or radiation exposure in cell lines.

Next, we used click-fluoro-quant to establish conditions to prevent tagging abundant background DNA modifications that can confound accurate calling of biologically relevant changes in the context of modification sequencing analysis. Namely, for mapping oxidized guanines, both FPG and ENDOIV are needed to generate 3′-OH groups as a marker of the oxidized-guanine site, however, 3′-OH groups exist independently. Furthermore, 3′-OH groups are created by the action of ENDOIV on pre-existing endogenous AP sites and phosphorylated breaks, and we wish to exclude these for accurate mapping of oxidized guanines. Therefore, we established a workflow that involved first ENDOIV treatment creating unspecific 3′-OH groups and afterwards the removal of the breaks by either repairing them with deoxynucleotide addition and ligation^52^ (Fig. 1g, repair) or blocking them by adding non-reactive dideoxynucleotides (ddNTP)^34^ to the sites (Fig. 1g, blocking). There were fewer unspecific 3′-OH groups after blocking vs repair (Fig. 1g), even after attempting to optimize the latter by increasing the incubation time and temperature as was done previously^53^ (Supplementary Fig. 1e), which informed us to adopt the blocking strategy for sequencing.

To address artefactual modifications, we evaluated their levels associated with gDNA-sample-handling factors such as purification method, pipette-tip-bore size, and heating (Supplementary Fig. 1f-g), which suggested avoiding heat inactivation of enzymes as causing an increase of modifications. In addition, we identified that the ddNTP blocking excluded most 3′-OH groups generated by sonication (Fig. 1h, sonication+blocking vs. sonication). Sonication is used to fragment gDNA to prepare libraries for sequencing and is well established as a major source for artifactual DNA modifications^42^. However, ddNTP-blocked non-sonicated gDNA still had almost 4-fold less unspecific 3′-OH groups than blocked sonicated samples (Fig. 1h, blocking vs. sonication+blocking). This informed us that, in order to prepare sequencing libraries, fragmentation by sonication should be performed after both ddNTP blocking of background 3′-OH groups (present or generated due to pre-existing breaks or AP sites) and after labeling of the target DNA modifications.

### Click-code-seq enables precise sequencing of oxidative DNA modifications in the human genome

Having established conditions to minimize background and artifactual DNA modifications with the assistance of click-fluoro-quant, we applied an optimized DNA-library-preparation protocol of click-code-seq, which had so far been used only to map oxidized guanines in yeast^34^ and a bacterial plasmid^37^, to now effectively map them across the human genome. Moreover, we upgraded click-code-seq by creating a conceptually new version of the code sequence called a Modified DNA Identifier Sequence (MoDIS). MoDIS is comprised of three modules, namely, a validation code (VC) of fixed nucleotide sequence, a randomized-index code (RIC), and an annealing-site sequence (AS) for specifically binding a primer for PCR amplification (Supplementary Fig. 2a). Due to the presence or absence of a VC in sequencing reads, we can distinguish sequencing reads that identify a DNA modification of interest from false-positive reads derived from amplification artefacts in library preparation (Supplementary Fig. 2b-d). Since MoDIS represents a mixture of sequences with different RIC, it allows for tagging PCR-amplified genomic sequences originating from different cells and genomes (Supplementary Fig. 2e-f).

We used MoDIS to generate single-nucleotide-resolution maps of endogenous DNA oxidation in gDNA from a human cancer cell line (Supplementary Fig. 2g-h). In detail, we isolated gDNA from HAP1 cells, treated it with ENDOIV to convert AP sites and phosphorylated breaks into 3’-OH breaks. These were ddNTP-blocked to conceal them from the oxidation map. Afterwards, we used FPG and ENDOIV to excise oxidized guanosines and Therminator IX to incorporate prop-dGTP into the resulting gaps. Only then was the DNA sonicated, end-repaired, and sequencing adapters were ligated. Next, we ligated biotinylated MoDIS to prop-dGTP sites via CuAAC, performed a streptavidin-based enrichment of DNA fragments ending at the original DNA-modification sites, PCR-amplified and sequenced the enriched fragments. With VC in MoDIS, we identified that around 15% of sequencing reads were unspecific and filtered them out (Supplementary Fig. 2d). Additionally, including RIC in MoDIS increased the number of identified oxidation sites by around 35% since without them identical reads originating from different cells/genomes would have been removed by deduplication (Supplementary Fig. 2f). Around 90% of identified DNA-modification sites were guanines (Fig. 2a), which reflected the guanine-oxidation products recognized by FPG, including 8-oxoG and fapy-G^54,55^, and indicated a high specificity of click-code-seq library preparation. The identified guanine-oxidation sites were supported by over 20 million unique reads (Supplementary Fig. 2g). Overall, the optimized protocol of click-code-seq including MoDIS and ddNTP blocking allowed highly specific sequencing of oxidative DNA modifications in the human genome.

**Fig. 2.**
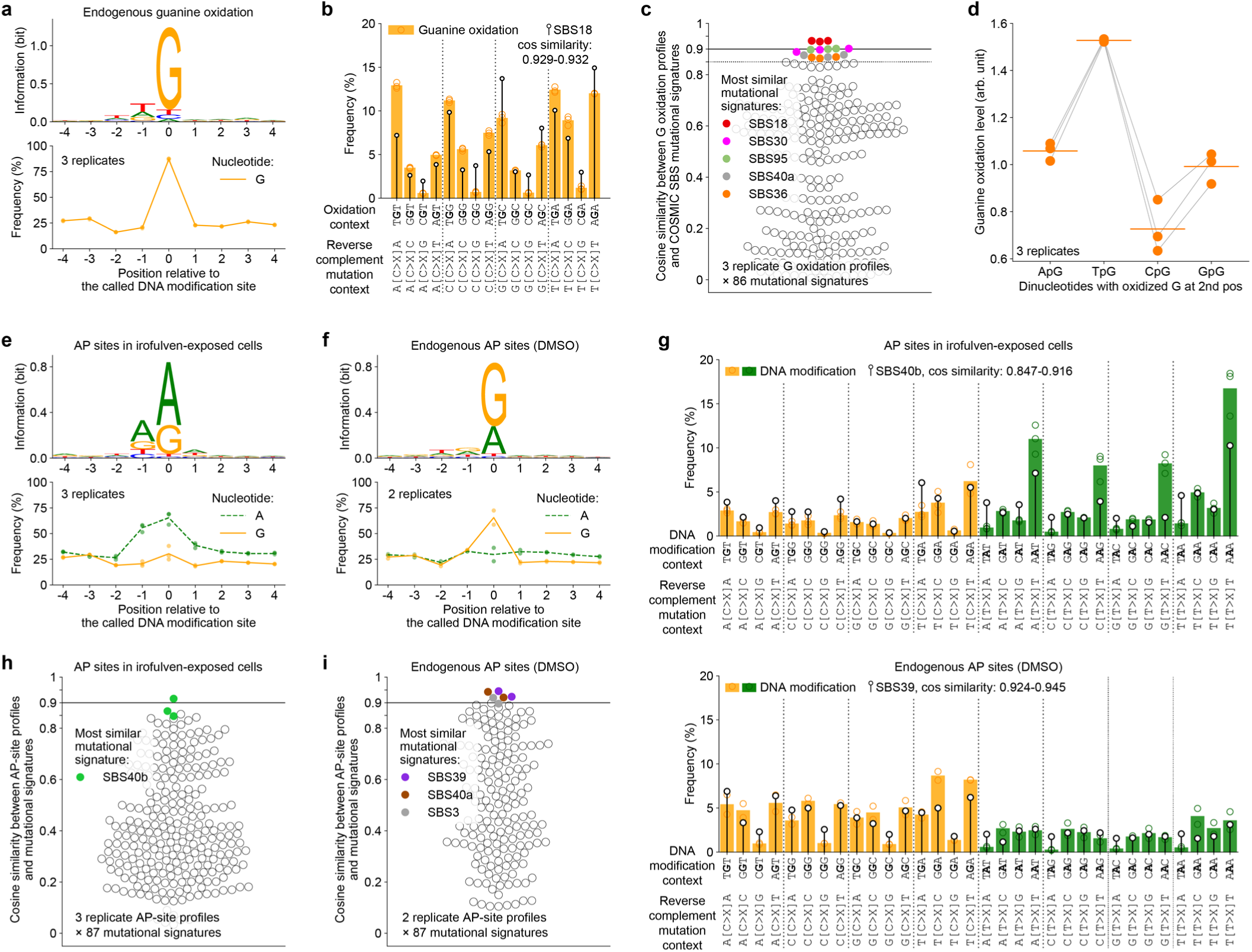
Click-code-seq uncovers single-nucleotide-resolution precursors of the ROS-related mutational signature SBS18 and identifies that irofulven induces depurination in adenine dimers. **a** Sequence logo and guanine frequency in the local genomic context around guanine-oxidation sites in HAP1 cells. **b** The distribution of oxidized guanines in trinucleotide contexts in HAP1 cells (n=3 replicates) matched with the ROS-related mutational signature SBS18 from human cancers (COSMIC). **c** The oxidation profile has the highest cosine similarity to SBS18 among all 86 COSMIC single base substitution signatures. **d** Guanine oxidation level is lower in CpG dinucleotide compared to other dinucleotides. Line-connected markers: biological replicates. Horizontal lines: mean. Arb. unit: **Methods** describe the normalization of DNA-modification level. **e-f** Sequence logo and frequencies of guanine and adenine in the local genomic context around AP sites in U2OS cells exposed to irofulven (**e**) or DMSO (**f**). **g** The distribution of AP sites regarding trinucleotide context in U2OS cells exposed to irofulven (upper, n=3) or DMSO (lower: n=2). The AP-site profiles are matched with the indicated mutational signature. **h-i** Comparison between the AP-site profiles from U2OS cells exposed to irofulven (**h**) or DMSO (**i**) and 86 mutational signatures from COSMIC plus one experimental illudin S mutational signature^56^. In panels **a,e,f**, sequence logo: merged data of all biological replicates; makers and line: frequency in replicates and the mean, respectively. In panels **b**,**g**, markers and filled bars: frequency of a trinucleotide with a mapped modification of G (**b**) or G and A (**g**) in the second position shown for each replicate and as the mean, respectively; lollipop: mutational signature shown for C substitution (87% of SBS18) in **b** or for C and T substitutions in **g**; frequencies of DNA modifications and mutations each add up to 100%; X: A, G and T considered together. In panels **c**, **h** and **i**, marker: cosine similarity between one mutational signature and one replicate DNA-modification profile, which is calculated by matching a vector of 32 trinucleotide-specific 5′N[C/T>X]N3′ substitution frequencies and a respective vector of DNA-modification frequencies (for guanine oxidation, values corresponding to T>X substitution are set to zero); N: one of 4 nucleotides; X: A, G and T considered together.

To assess the genomic-site specificity of guanine oxidation at the single-nucleotide level, we examined if the same or different guanines are found oxidized across replicate mapping experiments. Here, we identified that approximately 13,574,049-14,065,470, or ∼90%, oxidized-guanine sites are unique for each of three replicate experiments (Supplementary Fig. 3a). In contrast, 259,973, or 1.7%, sites had oxidized guanine consistently across all three replicates. These reproducibly detected sites were identified by more reads with individual RICs (mean: 5.3-7.3 across replicates, 90^th^ percentile: 10-13) compared to replicate-specific guanine-oxidation sites (mean: 1.3 across replicates, 90^th^ percentile: 2) (Supplementary Fig. 3b), reflecting that more cells had an oxidized guanine at a reproducibly detected site compared to a replicate-specific site. Despite about 90% of detected sites being unique to each experiment at the single-nucleotide level, aggregating the data into genomic bins showed the robustness of the oxidation maps, with pair-wise correlations of the 100-Kb-binned data above 0.88 among triplicate analyses (Supplementary Fig. 3c).

### DNA-oxidation profile in HAP1 cells correlates with cancer mutational signatures of ROS exposure

To understand how patterns of DNA oxidation in cultured human cells may relate with analogous mutational patterns from actual human cancer genomes, we determined the counts of oxidized guanines as a function of trinucleotide context (Fig. 2b) and compared this profile with single base substitution (SBS) signatures from the Catalogue of Somatic Mutations in Cancer (COSMIC)^57,58^ (Fig. 2c). Remarkably, the most similar SBS signature to the experimental data was SBS18, with cosine similarity in the range of 0.929-0.932 across replicate oxidized-guanine profiles (Fig. 2c). SBS18 was established to have ROS-induced etiology according to experiments with human induced pluripotent stem cells exposed to potassium bromate^59^. Above the cosine-similarity threshold of 0.85, two other putative ROS-related mutational signatures matched the experimental oxidation profile, namely: SBS30, associated with *NTHL1* deficiency^60^ (0.897-0.902 cosine similarity), and SBS36, associated with *MUTYH* deficiency^61,62^ (0.864-0.869 cosine similarity). Despite the overall high similarity between the damage profile and the mutational signature SBS18 (Fig. 2b, filled bars vs lollipops), the mutation frequency is markedly higher than the oxidation frequency at certain trinucleotides, such as TGC, AGA and all CpG dimers CGN, which may indicate impaired repair in those particular contexts. On the contrary, there is an almost two-fold drop in the frequency of mutations relative to oxidation in TGT and GGG trinucleotides, potentially reflecting preferential repair (Fig. 2b). By relating the numbers of oxidized guanines in different sequence contexts to the abundance of these contexts in the human genome, we observed that the probability of guanine to be or remain oxidized is the lowest if this guanine is preceded by cytosine (CpG sites) compared to other nucleotides (Fig. 2d). As CpG sites are substrates of cytosine methylation, this suggests a potential relationship of DNA oxidation and DNA methylation.

### Drug-induced depurination is elevated in adenine dimers

Chemical and drug-induced DNA alkylation can promote base hydrolysis, leading to the formation of abasic (predominantly apurinic, AP) sites. To understand the genome-wide patterns of alkylating-agent-induced AP sites, we analyzed gDNA from a human cancer cell line (U2OS) exposed to the chemotherapeutic drug irofulven (6-hydroxymethylacylfulvenein, HMAF; 15 µM)^50,51^. Irofulven-induced alkylation sites in extracted gDNA were allowed to convert to AP sites (37 °C, 18 h; spontaneous depurination half-life is in the range of 2-8.5 h^63^). We blocked DNA breaks by adding ddNTP with Therminator IX, converted all AP sites, whether drug-induced or endogenous, to gaps using ENDOIV. Next, we incorporated prop-dGTP and prop-dATP in the gaps, followed by further click-code-seq steps, the same as described above for oxidation mapping. By the identical procedure, we mapped endogenous AP sites in vehicle (0.03% DMSO) exposed cells. Additionally, by click-fluoro-quant analysis of samples under the same conditions, we observed around 5-fold more AP sites in the gDNA of irofulven-exposed cells compared to vehicle-exposed cells (Fig. 1f, 15 µM irofulven), suggesting there to be 5-fold more drug-induced than endogenous AP sites in the AP-site sequencing of irofulven-exposed cells.

Consistent with expectations, we identified mostly adenine and guanine at the position of the called DNA-modification sites in both irofulven- and vehicle-exposure conditions (Fig. 2e-f). Of note, the irofulven-induced depurination was observed more frequently at adenosines (Fig. 2e), in line with existing knowledge of acylfulvene DNA-alkylation chemistry^64^. However, endogenous depurination (*i.e.*, in vehicle-exposed cells) was mostly detected for guanosines (Fig. 2f), which agrees with snAP-seq^27^ but contradicts SSiNGLe-AP data^28^. For irofulven-exposed cells, we obtained 31,804,203-42,593,605 unique reads (Supplementary Fig. 2g) corresponding to 27,052,371-35,050,910 AP sites (Supplementary Fig. 3d). Over 90% of these AP sites were unique to each replicate but 283,400, or 0.8-1%, were found consistently across all three replicates (Supplementary Fig. 3d) and with more reads with individual RICs compared to replicate-specific AP sites (Supplementary Fig. 3e). Despite the low genomic-site specificity of depurination at single-nucleotide resolution, the AP-site maps in irofulven-exposed cells were highly similar when the data were aggregated in consecutive bins, with 100-Kb binning resulting in the correlations of 0.98 for A and 0.96 for G among the replicates (Supplementary Fig. 3f). Similar observations were made for endogenous AP sites (Supplementary Fig. g-i).

When using the same strategy to analyze how AP sites distribute amongst trinucleotide contexts, we found that the AP sites in irofulven-exposed cells were preferentially present at adenine dimers (Fig. 2g, upper half, trinucleotides AAT, AAG, AAC and AAA). The profile of endogenous depurination was drastically different, with the highest DNA modification levels at GGA and AGA trinucleotides (Fig. 2g, lower half). The DNA-modification pattern in irofulven-exposed cells was only moderately similar (cosine similarity 0.707-0.746) to the recently reported mutational signature of illudin S^56^, the natural-product precursor of irofulven (Supplementary Fig. 3j). A large discrepancy between both distributions is observed at TA dimers TAN (Supplementary Fig. 3j), suggesting a high mutagenicity of the relatively less frequent irofulven-DNA adduct in this sequence context, differences in sequence specificity of modification formation and repair between irofulven and illudin S, or other experimental differences between the studies. Comparing the DNA-modification pattern in irofulven-exposed cells to COSMIC mutational signatures, we surprisingly uncovered a strong match with SBS40b (cosine similarity 0.847-0.916; Fig. 2g, upper half; Fig. 2h), a recently discovered unknown-etiology signature strongly associated with clear cell renal cell carcinoma and biomarkers of impaired kidney function^65^. The pattern of endogenous AP sites was also highly similar to two mutational signatures of unknown etiology, namely, SBS39^66^ (cosine similarity 0.924-0.945; Fig. 2g, lower half; Fig 2i) and SBS40a^67^ (cosine similarity 0.921-0.943; Fig. 2i). Thus, the analysis of DNA-modification levels in the context of flanking nucleotides may be useful to understand the etiology of mutational signatures found in human cancers: SBS40b may be due to an exposure to an alkylating agent with similar biochemical properties to irofulven, and SBS39 and SBS40a may be due to endogenous depurination.

### DNA-oxidation and depurination landscape does not correlate with chromatin state

Moving from single-nucleotide level to large genomic bins, we noticed varying guanine-oxidation levels along chromosomes even after correcting for the guanine count per bin (1^st^ percentile 0.68-0.73 across replicates, 99^th^ percentile 2.15-2.22 at 100-Kbp binning, Supplementary Fig. 3k). To test if this variation can be explained by chromatin state, we matched the guanine-oxidation levels with chromatin features, derived from the ChIP-Atlas^68^ standardized re-analysis of published data for the same unexposed cell line (HAP1). Here, we found that, at 100-Kbp resolution, guanine-oxidation levels positively correlated with euchromatin features, namely, DNase I hypersensitivity sites (transcriptional activity), H3K4me1 (active enhancers), H3K4me3 (active gene promoters), H3K36me3 (active gene bodies) and H3K27ac (active promoters and enhancers) (Fig. 3a, 100 Kbp). There also was a positive, albeit much weaker, correlation with heterochromatic marks, namely, H3K27me3 and H3K9me3 (Fig. 3a, 100 Kbp). Since the average ChIP-Seq peak size for the chromatin marks is in the range of 118-1,023 bp (HAP1), we more finely binned the ChIP-Seq and guanine-oxidation data and observed that the correlations were not preserved at higher resolutions, falling to around zero at 1-Kbp binning (Fig. 3a). We made similar observations after matching irofulven-induced (Fig. 3b, Supplementary Fig. 3l) or endogenous (Fig. 3c, Supplementary Fig. 3m) AP sites with chromatin features in the U2OS-cell genome at increasing resolutions. Thus, while the presence of an eu- or heterochromatin mark in a genomic bin is weakly associated with higher DNA-modification levels at 100-Kbp resolution, the distributions of guanine oxidation and AP sites do not seem to be primarily shaped by chromatin state locally, *i.e.*, at 1-Kbp resolution, where other genomic features could have more influence.

**Fig. 3.**
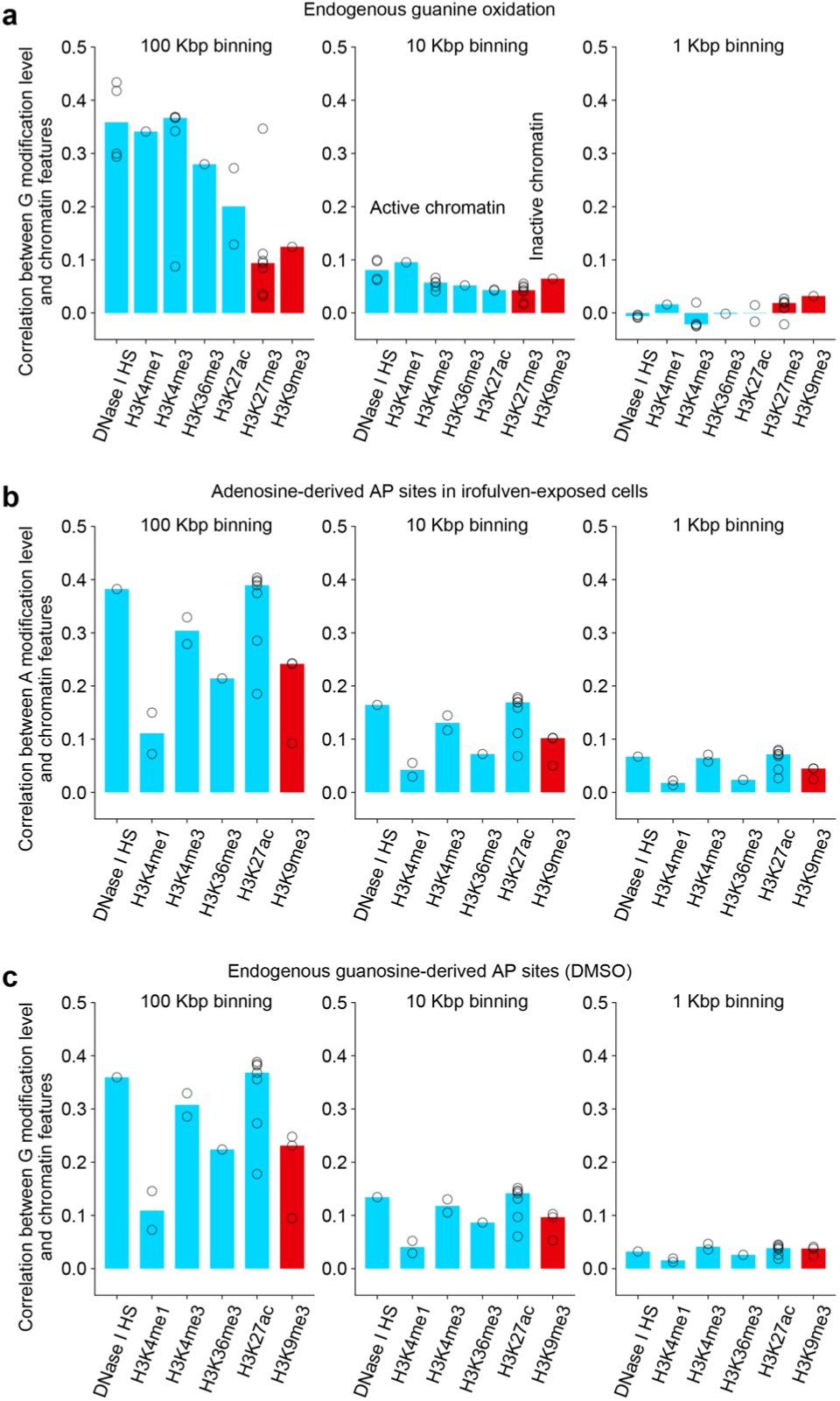
DNA-oxidation and depurination levels correlate with chromatin marks on large genomic bins, however, this association breaks down in more local genomic contexts. **a-c** Spearman correlation coefficient for the indicated chromatin marks and the level of endogenous guanine oxidation in HAP1 cells (**a**), irofulven-induced AP sites at adenines in gDNA from U2OS cells (**b**) or endogenous AP sites at guanines in gDNA from vehicle (DMSO)-exposed U2OS cells (**c**) with 100-, 10- or 1-Kbp genome binning. DNA-modification data: for each replicate experiment, signal was summed within bins spanning the genome, normalized by G (**a,c**) or A (**b**) counts per bin obtained from the reference genome and adjusted by replicate-specific normalization factor reflecting sequencing depth; n=3 (**a-b**) or n=2 (**c**) replicate data sets were averaged per bin and then used for the correlation. Chromatin marks: each marker indicates a ChIP-Seq mapping in a corresponding cell line under control conditions (without an exposure or genetic intervention) retrieved from ChIP-Atlas; binning of ChIP-Seq peak is described in **Methods**; a bar is median correlation coefficient across available ChIP-Seq experiments for a chromatin mark. Average ChIP-Seq peak sizes in HAP1 (**a**): 118-1,023 base pairs; in U2OS (**b-c**): 79-636 base pairs. Bin numbers: 28,601 in 100-Kbp binning; 285,360 for G modification and 285,354 for A modification in 10-Kbp binning; 2,852,127 for G modification and 2,852,122 for A modification in 1-Kbp binning. Kbp: 1,000 base pairs.

### DNA-oxidation levels are higher with increasing gene expression and have a strand bias

Exploring DNA-modification patterns in functional elements of the genome, we measured guanine-oxidation levels in genes between transcription start and end sites, separately in the transcribed and non-transcribed strands, and related these levels to gene expression. To account for different guanine abundances in the strands, we computed guanine-oxidation level as the ratio of number of mapped oxidized guanines to all guanines in the respective gene and strand. To compare guanine-oxidation levels amongst replicate samples independent of sequencing depth, we normalized all genomic-feature-specific guanine-oxidation levels by the median guanine-oxidation level in the transcribed (antisense) strands of unexpressed genes (Fig. 4a; the median of the second yellow box is set to 1 by this normalization). We found no overall strand bias for occurrence of guanine oxidation in the group of unexpressed genes (Fig. 4a, unexpr). However, with increasing gene expression, guanine-oxidation levels continuously increased, especially in the non-transcribed vs. transcribed strand (Fig. 4a). This gene-expression-dependent strand bias of guanine oxidation was robust across biological replicates, with median levels increasing maximally by 14.6% for the non-transcribed strand, however, only by 6.3%, for the transcribed strand, both relative to the unexpressed gene tier (Fig. 4b).

**Fig. 4.**
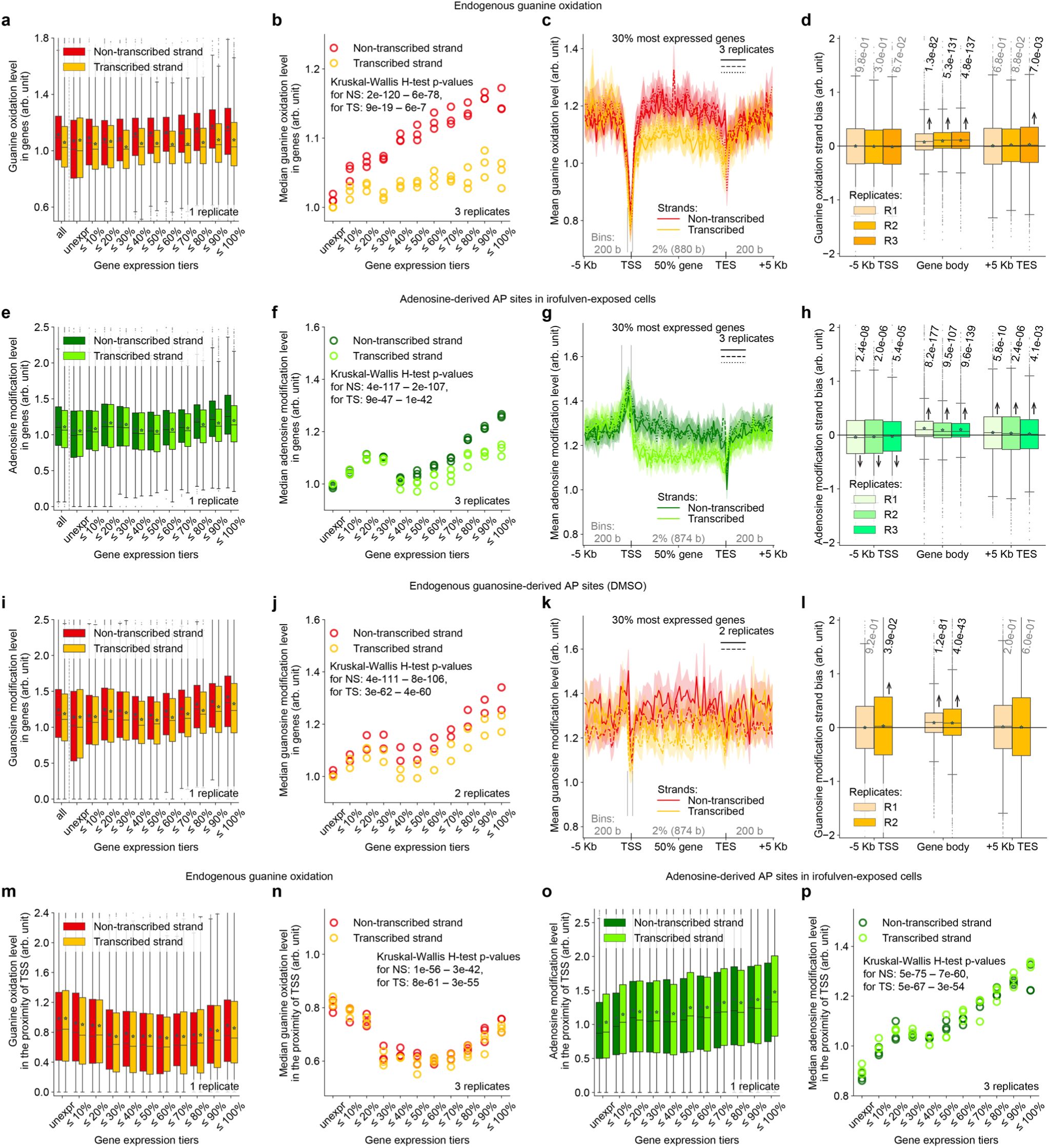
Oxidized guanines and AP sites accumulate in gene bodies with increasing gene expression, especially on the non-transcribed strand. **a-d, m-n** Endogenous guanine oxidation in gene bodies and adjacent regions in gDNA of HAP1 cells. **e-h, o-p** Irofulven-induced AP sites at adenines in gene bodies and adjacent regions in gDNA of U2OS cells. **i-l** Endogenous (DMSO) AP sites at guanines in gene bodies and adjacent regions in gDNA of U2OS cells. Panels **a,e,i**: DNA-modification level in each strand of the gene (from TSS to TES) body in protein-coding genes from one replicate experiment versus gene expression level; first, all genes are grouped together; after the dashed line, the genes are tiered according to expression level. Panels **b,f,j**: median DNA-modification level in each strand of the gene body across protein-coding genes in multiple replicates versus gene expression level. Panels **c,g,k**: strand-specific profile of the mean DNA-modification level and its 95% c.i. throughout the gene body and its upstream and downstream regions in highly expressed genes (**c**: n=3,994; **g,k**: n=4,425); solid, dashed, and dotted curves with shades: different biological replicates; genes are aligned at TSS and TES; vertical dashes demark TSS-adjacent regions analyzed in **m-p**. Panels **d,h,l**: Strand bias (difference) of DNA-modification levels between the non-transcribed and transcribed strands within the gene body and adjacent upstream and downstream 5-Kb regions for 30% most expressed genes (**d**: n=3,994; **h,l**: n=4,425) in multiple replicates; we provide p-values of two-tailed Wilcoxon signed-rank test; arrow shows the shift of the median from 0 if p-value<0.05. Panels **m-p**: DNA-modification level around the TSS (**m-n**: between -400 and +600 b; **o-p**: between -1000 and 0 b) in each strand in protein-coding genes versus gene expression level; plots are built analogously to **a-b**. In panels **a,d,e,h,i,l,m,o**, boxes are interquartile ranges, internal horizontal lines are medians, stars are means, whiskers extend to the furthest datapoint within 1.5x interquartile range, datapoints beyond are shown as small markers; number of genes per tier and number of genes beyond the minimal and maximal y-axis values are indicated in **Methods**. In panels **b,f,j,n,p**, we provide the min-max ranges of p-values of the Kruskal–Wallis H test performed for either strand across all replicates. Arb. unit: **Methods** describe DNA-modification level normalization. TTS and TES: transcription start and end sites. Kb: kilobase; b: base.

To confirm that the observed dependence of guanine oxidation on strand assignment and gene-expression level (Fig. 4a-b) is not due merely to confounders like varying sequence composition throughout gene expression tiers or different guanine oxidation propensities across trinucleotides (Fig. 2b), we analyzed the same relationship on the level of individual trinucleotides (Supplementary Fig. 4a). Here, we found that the median guanine-oxidation level changed with gene expression in 15 out of 16 trinucleotides. In 12 cases, we observed generally increasing dependencies with different maximal fold changes compared to the unexpressed-gene tier, namely, around 15.5-29.5% in AGA, TGN, AGG, GGA, AGC, 38.2% in GGG, 50.1% in AGT, 58.9-60.3% in GGC, GGT for the non-transcribed strand (Supplementary Fig. 4a). In CpG dimers CGG, CGC and CGA, median guanine-oxidation levels were zero in unexpressed and lowly expressed genes, then increased to a peak at around the 60% gene-expression tier and dropped to zero again in the most expressed genes (Supplementary Fig. 4a). Interestingly, we identified that the extent of the guanine-oxidation strand bias also varies across the trinucleotides, being most pronounced in GGT, GGA, TGT (17.6-21.6%), virtually absent in AGC, CpG dimers CGN, and intermediate in the rest (Supplementary Fig. 4a). Thus, the overall guanine-oxidation level and its strand bias increase as function of gene expression, with most individual trinucleotides within genes showing the same pattern, however, to varying degrees.

For genomic locations of highly expressed genes, we observed a strand bias of guanine oxidation only throughout the gene body but not upstream of the transcription start site (TSS) or downstream of the transcription end site (TES) (Fig. 4c, Supplementary Fig. 4b; compare with analogous profiles of unexpressed genes: Supplementary Fig. 4c-d), which was further supported by Wilcoxon signed-rank test (Fig. 4d). The low oxidation levels in the transcribed strand may be evidence that oxidized guanines, or their base-excision-repair (BER) intermediates are repaired in a transcription-coupled (TC) manner. These results are consistent with the findings of the involvement of nucleotide excision repair (NER) in the removal of 8-oxoG^69^ but contradictory to an earlier observation of no strand bias in the repair of a housekeeping gene^70^.

### AP sites accumulate with increasing gene expression and have a strand bias

To explore how transcription may relate with irofulven-induced and endogenous AP sites, we analyzed their gene-expression dependency, separately for adenosines and guanosines, presenting the gene- and strand-specific data of one biological replicate (Fig. 4e,i, Supplementary Fig. 5a,e) and medians of multiple replicate experiments (Fig. 4f,j, Supplementary Fig. 5b,f). For irofulven-induced AP sites derived from adenosine (Fig. 4f) and guanosine (Supplementary Fig. 5b), we observed, respectively, 27.3% and 26.3% elevations of median DNA modification level for the non-transcribed strand in the tier of the most expressed genes (≤100%) compared to unexpressed ones. For the transcribed strand, the maximal elevation of median DNA modification was 13.1% for adenosines and 11.4% for guanosines (Fig. 4f, Supplementary Fig. 5b), creating a pronounced strand bias of the DNA modifications throughout the second half of gene-expression tiers (maximally 11.5% for both A and G), which is in line with irofulven-induced adducts being known TC-NER substrates^71,72^.

We identified that irofulven-induced AP sites derived from adenosine alkylation prevail in the non-transcribed strand throughout the gene body and downstream of TES (Fig. 4g), as reproduced in three replicates (Fig. 4h), however, the strand bias is swapped upstream of TSS such that the AP sites occur more frequently in the transcribed strand (Fig. 4g-h). This swapping-strand-bias signature was recently observed for another TC-NER-specific substrate, a DNA alkylation product of the chemotherapeutic drug trabectedin^15^, and may be related to divergent transcription from promoters^73^. A significant strand bias of irofulven-induced AP sites derived from guanosine was found only throughout the gene body (Supplementary Fig. 5c-d). As for endogenous depurination, we also observed elevating AP-site levels (maximally 49.6% for A, 27.7% for G) and a widening strand bias with increasing gene expression (maximally 10% for A, 8% for G), only within the gene body, both for guanosines (Fig. 4i-l) and adenosines (Supplementary Fig. 5e-h). We observed that DNA-modification burdens increased with gene expression for almost all trinucleotides in both irofulven-induced and endogenous depurination, yet trinucleotides varied in maximal fold changes and strand biases of AP-site levels (Supplementary Fig. 6 and 7). Particularly for adenosine-derived AP sites, gene-expression-dependent patterns were notably different between vehicle- and irofulven-exposed samples, supporting that the endogenous adenosine-derived AP sites do not confound the mapping of irofulven-induced ones (Supplementary Note 1).

Overall, by establishing the strand bias of DNA-modification levels as a signature of transcription-coupled repair via the case of depurination induced by a known TC-NER-specific drug irofulven (Fig. 4e-h), the presented data may also suggest transcription-coupled repair of endogenous DNA oxidation (Fig. 4a-d) and depurination products (Fig. 4i-l and Supplementary Fig. 5e-h).

### DNA oxidation and depurination diverge in the proximity of transcription start site

Besides the observations of strand bias described above, another prominent feature of the DNA-modification profiles with respect to gene regions is a signal deviation in the proximity of TSS. For example, close to TSS, there are a sharp drop of guanine oxidation (Fig. 4c) (which has been disputed in literature with both depletion^29,34,35^ and enrichment observations^30,38,39^) and a spike of irofulven-induced adenosine depurination (Fig. 4g). To further explore these features, we measured guanine-oxidation level in the signal drop between -400 and +600 bases around TSS (vertical dashes in Fig. 4c) and related these values to gene expression. Surprisingly, we discovered a non-monotonic relationship (Fig. 4m-n) in which guanine oxidation is highest in unexpressed genes, decreases and reaches a minimum (25.4-29.2% reduction) in the 50% and 60% tiers of gene expression, but then increases again (by around 23%). Nonetheless, guanine-oxidation levels in highly expressed genes were still lower than in unexpressed genes (Fig. 4m-n). The levels of adenosine-derived AP sites, both irofulven-induced and endogenous, in the proximity of TSS (-1000 and 0 bases, vertical dashes in Fig. 4g and Supplementary Fig. 5g) had a different behavior (Fig. 4o-p, Supplementary Fig. 8a-b), almost always increasing as a function gene expression with a maximal fold change of 45%-46.8% (Fig. 4p, Supplementary Fig. 8b) and different from the patterns of guanosine depurination (Supplementary Fig. 8c-f). Thus, burdens of DNA oxidation and alkylation-induced and endogenous depurination within the promoter regions of genes vary in relation to gene expression level, however, the respective associations have different forms, which may reflect different potential influences of these modifications or their repair on gene-expression regulation.

### Strand asymmetry of endogenous guanine oxidation and adenosine depurination in mtDNA

We analyzed modifications in mtDNA, which is particularly vulnerable due to its proximity to the electron transport chain, a major source of reactive oxygen species (ROS), and lack of histone protection^74^. The heavy (**–**) and light (**+**) strands of mtDNA differ in guanine content (Supplementary Fig. 9a, dashed bars). Interestingly, we observed that oxidized guanines were more prevalent in the – strand than expected based on its guanine content (Supplementary Fig. 9a, filled vs dashed bars). To characterize the propensity for oxidized guanine to exist throughout mtDNA, we aggregated oxidized guanines in consecutive 100-base bins for each strand. We normalized the bin- and strand-specific oxidized-guanine count to the total oxidized-guanine count in mtDNA and further to the bin- and strand-specific guanine count from the reference genome (Fig. 5a, left). Two prominent peaks were found in the **–** strand, specifically in the transcribed strand of the gene *ATP6* encoding ATP synthase membrane subunit 6 (peak bin: 8,600-8,700 bp, gene: 8,526-9,206 bp; 0-based coordinates) and in the D-loop region (peak bin: 16,100-16,200 bp, D-loop region/non-coding region: 16,023-16,568;0-575). Overall, most of the bins in the **–** strand had higher guanine-oxidation levels compared to their counterparts in the **+** strand, a bias confirmed by Wilcoxon signed-rank test (Fig 5a, right). Notably, since the transcribed strand of most mtDNA genes is the **–** strand, this bias contrasts with our findings in the nuclear genome, where transcribed strands typically showed lower guanine oxidation (Fig. 4a-d).

**Fig. 5.**
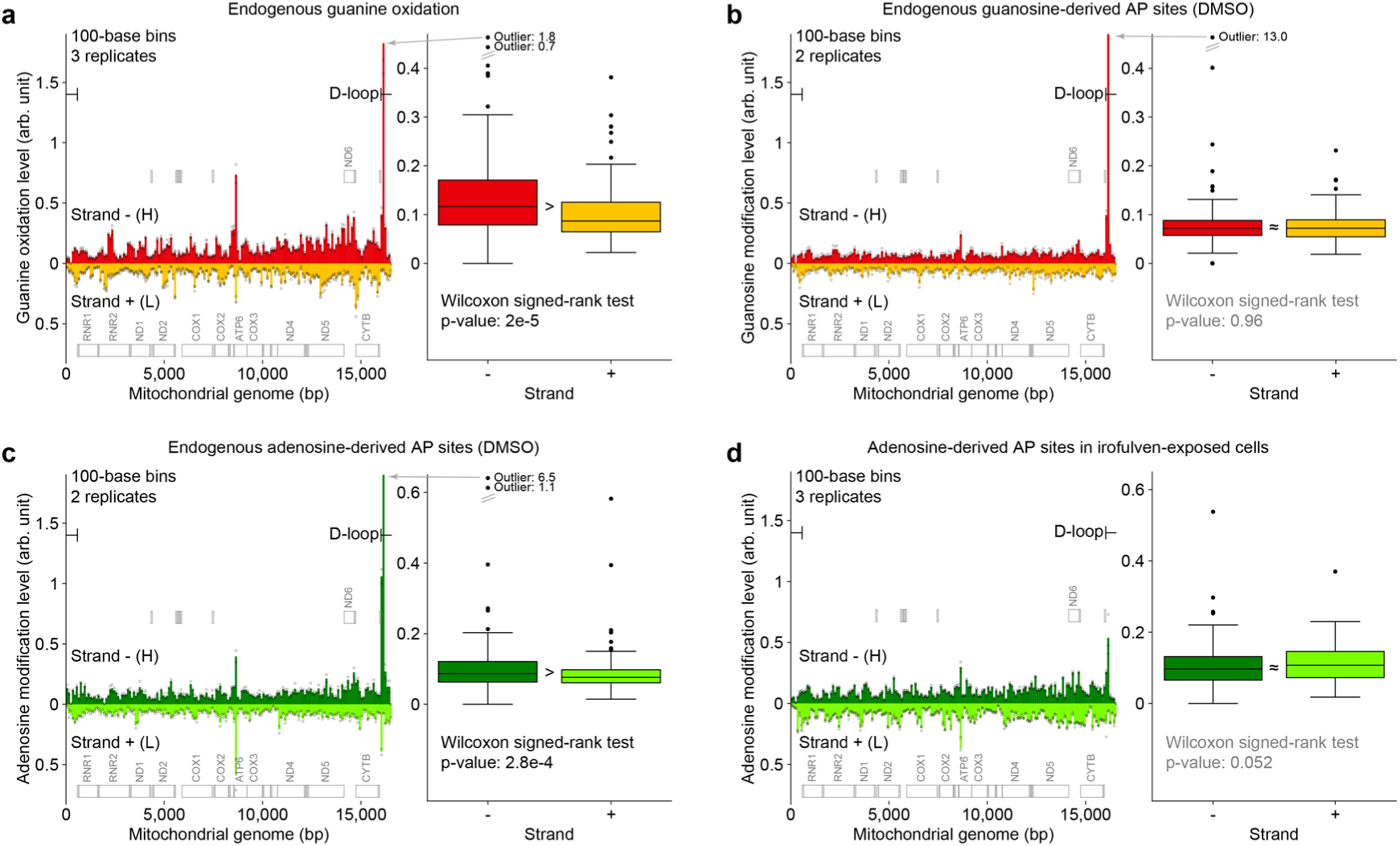
Endogenous guanine oxidation and adenosine depurination are more frequent in the – (heavy) strand of mtDNA, while endogenous guanosine depurination and drug-induced depurination do not have an overall strand bias. **a-d** Left panel: The strand-specific profile of endogenous guanine oxidation in HAP1 cells (**a**), endogenous guanosine (**b**) and adenosine (**c**) depurination in U2OS cells (DMSO vehicle), and adenosine depurination in irofulven-exposed U2OS cells (**d**) throughout mtDNA. Markers: replicates (number specified); bars: means of replicate values. The locations of 37 genes are provided as gray boxes along their sense strands. H: heavy; L: light. Right panel: bin-specific mean values from the left panel aggregated within each strand. Wilcoxon signed-rank test: 166 pairs of strand-specific values, two-sided alternative. Boxes are interquartile ranges, internal horizontal lines are medians, whiskers extend to the furthest datapoint within 1.5x interquartile range, datapoints beyond are shown as markers, the values of the most extreme outliers are provided. Arb. unit: **Methods** describe DNA-modification level normalization.

By a similar analysis, endogenous guanosine-derived AP sites showed no significant overall strand bias (Fig. 5b), while the less abundant (Supplementary Fig. 9b) endogenous adenosine-derived AP sites mapped in the same samples were significantly more frequent in the **–** strand than in the **+** strand throughout mtDNA (Fig. 5c), consistent with recent findings in mouse tissues^28^. Upon irofulven exposure, most AP sites originated from adenosine (Supplementary Fig. 9c), with no significant strand bias for both adenosine- and guanosine-derived AP-sites (Fig. 5d, Supplementary Fig. 9d).

Examining single-nucleotide coverage by reads with unique RICs, a proxy for frequency in a cell population, we revealed that guanine at position 16,103 (G16,103|**–**) and adenine at position 16,104 (A16,104|**–**) in the **–** strand had the highest modification coverage throughout mtDNA in respective mapping of oxidized guanines, guanosine- and adenosine-derived AP sites (Supplementary Fig. 9e-i). These single-nucleotide signals were the largest contributors to the D-loop-localized peaks observed in the bin analyses (Fig. 5a-d, Supplementary Fig. 9d). The D-loop region includes a third strand, 7S DNA, which represents a linear copy of the **–** strand of unknown function^75^ with the 3′-end nucleotide predominantly at position 16,105 and less abundantly at positions 16,104 and 16,103^76^. The uncovered DNA-modification signals at G16,103|**–** and A16,104|**–** may thus be localized in either 7S DNA or in the circular **–** strand. The occurrence of such strong DNA-modification signals close to the 3′ end of 7S DNA may suggest their functional relevance in the control of 7S DNA synthesis termination and hence mtDNA replication^75,77^. Thus, mtDNA exhibits an asymmetric distribution of endogenous guanine oxidation and adenosine depurination and produces strong DNA-modification signals at two single-nucleotide positions in the D-loop region.

## Discussion

In this study, we uncovered new insights into the non-homogeneous distribution of endogenous DNA oxidation and AP sites in human genomic DNA and created the first genome-wide map of DNA modifications induced by irofulven, an emerging precision-oncology chemotherapeutic drug^45,46^. This was achieved through the development and application of two complementary methods: click-fluoro-quant, a rapid and cost-effective fluorescence-based assay for quantifying oxidized guanines, AP sites, and DNA breaks; and click-code-seq, a genome-wide single-nucleotide-level DNA-modification sequencing assay. The endogenous guanine-oxidation profile in human cells was found to closely match the ROS-associated mutational signature SBS18 observed in cancers. Notably, CpG dimers were the least oxidized among guanine-containing dinucleotides, suggesting an interplay between DNA oxidation and methylation. The AP-site profile in irofulven-exposed cells was enriched in adenine dimers and was similar to the kidney-cancer-associated mutational signature SBS40b. Moreover, we discovered that the transcribed strands of genes suffer less from oxidized guanines and AP sites than the non-transcribed strands, with this strand bias widening with increasing gene expression. Finally, the data revealed that also mtDNA strands have unequal levels of endogenous guanine oxidation and adenosine depurination.

In this work, we addressed several basic experimental factors that can decrease the specificity of DNA-modification sequencing. Firstly, target-versus-background-differentiation issues generally affect DNA-modification sequencing methods such as in SSiNGLe-AP^28^ (AP sites vs breaks), CLAPS-seq^29^ (8-oxoG vs polymerase-stalling modifications like AP sites), GLOE-Seq^21^ and nick-seq^78^ (enzymatically derived breaks vs pre-existing breaks) and 250-bp-resolution AP-Seq^35^ (8-oxoG vs AP sites). Since click-code-seq relies on tagging available 3’-OH groups, background DNA breaks and AP sites can confound the mapping of oxidized guanines recognized through glycosylase and endonuclease treatment; similarly, background DNA breaks can confound AP-site mapping. We used click-fluoro-quant data to assess the efficiency of library preparation procedures to mitigate this, and blocked background modifications, as has been done in SSiNGLe-AP^28^ and nick-seq^78^. Secondly, for DNA fragmentation, we postponed sonication until after labeling target modifications since sonication is well known to produce unintended DNA modifications, such as 8-oxoG, leading to false positives in sequencing data^42^. The over-100-bp-resolution methods like OG-Seq^30^ and OxiDIP-Seq^39^ involve DNA sonication prior to labeling 8-oxoG; the issue was addressed for those methods by the addition of antioxidants. In AP-site-specific SSiNGLe-AP^28^ and thymidine-glycol-specific DPC-Seq^79^, DNA was fragmented enzymatically rather than through sonication. Thirdly, most DNA-modification sequencing methods rely on PCR amplification of enriched DNA fragments, potentially generating unspecific products due to spurious priming and making it impossible to distinguish whether a DNA-modification site occurs in one or multiple cells. We devised a new sequencing adapter, MoDIS, comprising a validation code (VC) to filter out unspecific PCR products and a randomized index code (RIC) to assess how frequently a genomic site is modified across a cell population. We thereby identified that approximately 15% of reads are unspecific, suggesting that spurious amplification products may significantly confound data from other DNA-modification methods lacking such an artefact control. Furthermore, the click-chemistry-aided ligation of MoDIS to DNA-modification sites may be more efficient than enzymatic ligation^80^ used in other sequencing methods, and the resulting biocompatible triazole linkage does not stall the polymerase during amplification^34,81^.

We found that, in human cells, both endogenous guanine oxidation (Fig. 4a-b) and irofulven-induced (Fig. 4e-f) and endogenous depurination (Fig. 4i-j) intensify in gene bodies with increasing gene expression, which aligns with a similar recent observation of endogenous AP sites in mouse tissues^28^. These findings suggest that DNA becomes more vulnerable to damaging agents, such as ROS, irofulven, and to hydrolysis during transcription. This vulnerability may be due to the single-strandedness of DNA during transcription, particularly, in transcription bubbles^82^ or potentially in co-transcriptional R-loops^83^. Single-stranded DNA provides greater accessibility of nucleobases to damaging agents, supported by higher rates of spontaneous depurination in single-stranded versus double-stranded DNA^44^. Transcription bubbles and R-loops could also explain the observed gene-expression-dependent strand bias (*i.e.* fewer DNA modifications in the transcribed strand of the gene body, Fig. 4d,h,l) as the transcribed strand is involved in DNA-RNA hybrid duplexes^82,83^, leaving the non-transcribed strand exposed. Alternatively, this strand bias could be attributed to transcription-coupled repair, which is initiated by RNA polymerase stalling and operates in the transcribed strand. The depletion of irofulven-induced AP sites in the transcribed strand supports this, as irofulven-DNA or, in general, acylfulvene-DNA adducts are known TC-NER substrates^64,72,84,85^. Similar strand biases for endogenous guanine oxidation and AP sites suggest that these modifications are also removed by transcription-coupled repair. Supporting this, AP sites have been shown to block transcription^86^ and are processed by TC-NER in yeast^87^. There is also evidence that NER contributes significantly to the removal of oxidative DNA modifications, where BER intermediates block transcription and are then resolved by TC-NER^69^. Interestingly, endogenous single-strand breaks also show a gene-expression-dependent strand bias, but contrary to oxidation and depurination, more breaks are formed in the transcribed strand^15^. These findings highlight the importance of transcription in the accumulation of different forms of DNA modifications in gene bodies.

Promoter regions are another genomic feature exhibiting intricate patterns of DNA modifications. For example, the strand bias of irofulven-induced adenosine-derived AP sites was inverted for at least 5 Kb upstream of the TSS, with the transcribed strand (from the gene-body perspective) accumulating more DNA modifications (Fig. 4g-h). This finding aligns with recent mapping of DNA breaks induced by trabectedin, another TC-NER-specific chemotherapeutic drug,^15^ and suggests TC-NER activity extends kilobases upstream of the TSS, likely connected to divergent transcription from promoters^73^. Interestingly, in line with the trabectedin study^15^, there is a 1-Kb region immediately upstream of the TSS that appears to be a TC-NER blind spot, where the level of irofulven-induced adenosine-derived AP sites peaks without a strand bias (Fig. 4g) and increases with gene expression (Fig. 4o-p), which may promote drug toxicity in a TC-NER-independent manner. Conversely, guanine oxidation in promoter regions sharply drops between -400 and +600 bp around the TSS (Fig. 4c), consistent with some studies^29,34,35^ but contradictory to others^30,38,39^. We found that promoter-region-localized guanine oxidation was highest in genes that are either highly expressed or not expressed (Fig. 4m-n). It is known that 8-oxoG in promoter regions can regulate gene expression via BER and altered G-quadruplex structures, as seen for *VEGF* and *NTHL1*^16,17^. Additionally, a recent genome-wide analysis reported reduced 8-oxoG in G-quadruplex sites but increased 8-oxoG in non-G-quadruplex potential-G-quadruplex sequences^29^. Our observations of varying levels of adenosine depurination and guanine oxidation in promoters as a function of gene expression may indicate a contribution of these modifications or their repair intermediates to transcriptional regulation.

While guanine oxidation and depurination mostly lacked genomic-site specificity at the single-nucleotide level (Supplementary Fig. 3a,d,g), we observed a high level of reproducibility when the data were aggregated at various levels, such as bins of 100 Kbp, 200 and 100 bases, functional elements, or trinucleotides. In line with the occurrence of AP-site hotspots in mouse tissues^28^, over 250,000 single-nucleotide sites were still reproducibly mapped across all replicate experiments for each DNA-modification type (Supplementary Fig. 3a,d,g). Two such signals occurred at G16,103|**–** and A16,104|**–** located in the D-loop region of mtDNA close to the 3′ end of the 7S DNA, where we observed the highest numbers of supporting reads (Supplementary Fig. 9e-i), which is potentially related to structures impacting 7S DNA synthesis termination and therefore mtDNA replication^75,77^.

Also in mtDNA, we observed that endogenous guanine oxidation and adenosine depurination are more prevalent throughout the heavy (**–**) strand, which is mostly the transcribed strand, compared to the light (**+**) strand (Fig. 5a,c). This contrasts patterns in the nuclear genome, where there is less guanine oxidation and depurination in the transcribed strand. This difference can be attributed to the widely accepted strand displacement model of mtDNA^77^, where heavy (**–**) strand synthesis begins significantly earlier than the light (**+**) strand, leaving the old heavy (**–**) strand unpaired and exposed. However, there is no such strand asymmetry for endogenous guanosine depurination (Fig. 5b) nor irofulven-induced depurination (Fig. 5d). We conjecture that their lack of asymmetry results from the opposing effects of mtDNA replication, which exposes the heavy (**–**) strand, and mtDNA transcription, which exposes the non-transcribed light (**+**) strand for most genes. In the cases of endogenous guanine oxidation and adenosine depurination, mtDNA replication may dominate, leading to the observed strand asymmetries. Such imbalance in DNA-modification distributions could be a basis of mechanisms driving mtDNA instability^88^.

The single-nucleotide nature of click-code-seq allowed us to correlate the frequency of DNA-modifications within trinucleotide contexts throughout the genome with mutational signatures found in human cancers, providing clues for the etiology of these signatures. Indeed, the highest similarity of the trinucleotide-specific pattern of endogenous guanine oxidation among all COSMIC mutational signatures was SBS18 (Fig. 2c), which is attributed to reactive oxygen species, further supporting this annotation. This observation provides an additional new example of forecasting human mutational signatures with DNA-modification signatures, such as observed for human lung cancers vs. cultured lung cells exposed to a tobacco carcinogen^32^ as well as cancers in patients previously treated with the glioblastoma drug temozolomide vs. cultured glioblastoma cells exposed to temozolomide^33^. In the present case, we uncovered a close match between the AP-site pattern in irofulven-exposed cells and the signature SBS40b, which is strongly associated with kidney cancer and seems to result from an unknown mutagenic exposure varying across countries^65^, indicating the exposure may be related to an alkylating agent with similar chemical properties to irofulven. Furthermore, a potentially relevant corollary in this case is the mutational signature attributed to aristolochic-acid exposure and kidney disease^89–91^, particularly given that the DNA adducts of aristolochic acid, like those of irofulven, have biological effects associated with TC-NER^92^.

In conclusion, we introduced here click-fluoro-quant for rapid, cost-effective quantitation of oxidized guanines, AP sites, and DNA breaks, and upgraded click-code-seq for single-nucleotide-precision mapping of oxidized guanines and AP sites in the human genome. We uncovered a non-uniform distribution of these DNA modifications across various genomic features and matched it with patterns of gene expression, DNA repair, replication and mutations. These tools and insights will advance research by characterizing processes that underlie genome integrity and function, with the potential to enhance precision medicine diagnostics and toxicology testing.

## Methods

Unless otherwise stated, chemicals were acquired from Sigma Aldrich, molecular biology reagents from New England Biolabs (NEB), and cell culture reagents from Thermo Fisher. Oligonucleotide DNA was synthesized and PAGE-purified by Eurogentec. Adapters and primers for click-code-seq were synthesized and HPLC-purified by Eurogentec. Solutions were prepared in MilliQ-purified H2O. Buffer EB acquired by Qiagen contains 10 mM TRIS-Cl, pH 8.5. Cell lines were authenticated by Microsynth AG, Switzerland.

### Cell culture, KBrO3, irofulven and UVA exposure

HAP1 cells (RRID:CVCL_Y019, Horizon Discovery) were cultivated in Iscove’s Modified Dulbecco’s Medium supplemented with 10% fetal bovine serum (Gibco) and 1% penicillin-streptomycin (Gibco) at 37°C, 5% CO2, 3% O2. U2OS cells (RRID:CVCL_0042) were cultivated in McCoy‘s general cell culture medium, modified with high glucose, L-glutamate, bacto-peptone and phenol red, and lacking sodium pyruvate and HEPES (4-(2-hydroxyethyl)-1-piperazineethanesulfonic acid) (McCoy‘s 5A, Gibco 16600082), supplemented with 10% fetal bovine serum (Gibco) and 1% penicillin-streptomycin (Gibco) at 37 °C, 5% CO2. BJ-5ta cells (ATCC) were cultured at 37 °C, 5% CO2, 3% O2 in a medium composed of four parts of Dulbecco’s Modified Eagle’s Medium (DMEM 1x + GlutaMAX, 31966-021, Gibco), one part of Medium 199 (22340-020, Gibco), 10% fetal bovine serum (FBS, SH30066.03HI, GE Healthcare), and 0.01 mg/mL Hygromycin B (SH30066.03HI, Sigma).

For KBrO3 treatment, medium was removed, and cells were washed with 10 mL Dulbecco’s phosphate-buffered saline (DPBS). KBrO3 was dissolved in 10 mL medium for a final concentration of 50 mM and added to the cells followed by a 30 min incubation at 37 °C and harvesting. For irofulven treatment, medium was prepared by adding HMAF (50 mM stock, dissolved in DMSO) to 10 mL medium to a final concentration of 0.8-15 μM. A control medium was prepared by adding DMSO in the same volume as the treatment to 10 mL medium. The treatment and control media were filtered with a sterile syringe filter (0.22 μm) and added to the cells followed by 4-h incubation at 37°C. For UVA treatment, medium was replaced with serum-free medium. Dishes were covered with soda-lime glass lids to filter our potential UVB contamination from the UV lamp. Cells were irradiated in a UV irradiation chamber BS-02 (Opsytec Dr. Gröbel, Germany) with an intensity of 8 mW/cm^2^ for a final dose of 10 J/cm^2^ (around 18 min) controlled by a dosimeter.

### gDNA extraction and sonication

Cells were harvested by removing the medium, washing once with 10 mL DPBS followed by addition of 1 mL 0.25% Trypsin EDTA solution (Gibco). After an incubation of 5 min at 37°C, 9 mL medium was added, cells resuspended and transferred to a Falcon centrifugal tube. Cells were centrifuged (4°C, 0.3 rcf, 5 min) and the supernatant removed. The cell pellet was either stored at -20°C or directly used for DNA extraction. gDNA was extracted from cell pellets using the Monarch Genomic DNA Extraction Kit, following the manufacturer’s instructions. In case of oxidative-modification analysis, DPBS used in the cell pellet resuspension step was supplemented with 1 mM deferoxamine and 50 µM *n-tert*-butyl-α-phenylnitrone to avoid artefactual oxidation. Extracted gDNA was quantified using the Quantus™ Fluorometer with QuantiFluor® ONE dsDNA Dye. Sonication of gDNA was performed using a Q800R2 sonicator (QSonica) with the following settings: 4°C, 20% amplitude, pulse 15 s on, 5 s off, total 5 min. The resulting fragments were analyzed on an Agilent 2200 Tapestation using a High Sensitivity D100 ScreenTape.

### Fluorescence labeling of DNA modifications in oligonucleotides

For 8-oxoG site labeling, pre-annealed oligonucleotide DNA (IL1-T30, IL-1_oxoG; Supplementary Table 1) was used in concentrations of 2, 1, 0.75 and 0.5 µM. For the negative control, 2 µM pre-annealed oligonucleotide DNA (IL1-T30, IL-1; Supplementary Table 1) was used. DNA was mixed with 1 µL FPG (NEB, 8 U/µL) and 1 µL T4 PNK (NEB, 10 U/µL) in 1x NEBuffer 2 (NEB) in a final volume of 10 µL. The reaction mixture was incubated for 1 h at 37°C directly followed by the addition of 1.5 µL prop-dGTP (Jena Biosciences, 2.5 mM), 0.5 µL NEBuffer 2 (NEB), 2.8 µL H2O and 0.2 µL Therminator IX (NEB, 10 U/µL) and incubated for 10 min at 60°C. DNA was purified using the Monarch PCR & DNA Cleanup Kit according to the manufacturer’s instructions using the oligonucleotide protocol and an elution volume of 6 µL. The CuAAC reaction was performed adding 4 µL AF594 picolyl azide (Jena Biosciences, 10 mM stock in DMSO), 4 µL DMSO, 2 µL premixed CuSO4:THPTA (THPTA from Lumiprobe; 20 mM Cu^2+^ and 200 mM THPTA)^93^ and 2 µL sodium phosphate buffer (1 M, pH 7.0). The reaction was started by adding 2 µL freshly prepared sodium ascorbate (400 mM) and incubated for 60 min at 37°C. The reaction was purified using the Monarch PCR & DNA Cleanup Kit with 20 µL for elution. 8 µL of each sample was used for analysis by 20% (w/v) denaturing polyacrylamide gel electrophoresis, performed at 230 V for 1 h. After imaging using a ChemiDoc XRS+ System (Bio-Rad), gels were stained with 1x GelRed (Biotium) for 10 min and imaged again. For fluorescence analysis, the remaining oligonucleotide DNA was mixed with 600 µL PBS. The sample was split in 3x 200 µL and pipetted in a non-clear 96 black well plate. For blank measurement, 3x 200 µL PBS were added to the plate. Fluorescence was recorded with an Infinite Pro M200 Plate Reader (Tecan) using the following settings: λEx = 580 nm, λEm = 610 nm, gain = optimal (or kept identical for replicates), 25 flashes, 20 µs integration time, 25°C.

### Blocking and repair of DNA modifications

4 µg gDNA was used for each sample. To minimize artefactual damage, samples were inverted or flicked instead of pipette mixing. Pronex Magnetic Bead purification was performed by adding 80 µL beads to a sample of 50 µL (adjusted with H2O), then following the manufacturer’s instructions and eluting with Buffer EB. For ddNTP blocking, gDNA was first incubated with 1 µL ENDOIV (NEB, 10 U/µL) in 1x ThermoPol Buffer (NEB) in a final volume of 20 µL for 20 min at 37°C. Then, 2.5 µL ddNTP mix (Jena Biosciences, 2.5 mM), 0.5 µL 10x ThermoPol Buffer, 1.8 µL H2O and 0.2 µL Therminator IX (NEB, 10 U/µL) was added followed by a 10 min incubation at 60°C. The repair mix modified from Zatopek *et al.* comprised gDNA with 1 mM NAD^+^ (NEB), 1 µL dNTP mix (NEB, 10 mM), 1 µL ENDOIV (NEB, 10 U/µL), 5 µL TAQ Ligase (NEB, 40 U/µL) and 0.5 µL BST DNA Polymerase FL (NEB, 5 U/µL) and 2 µL 10x ThermoPol Buffer (NEB) in a total volume of 20 µL and was incubated for 30 min or overnight at 37°C. The repair mix from Shu *et al.* contained 2 μL ENDOIV (NEB, 10 U/µL), 1 μL BST DNA Polymerase FL (NEB, 5 U/µL), 2μL TAQ Ligase (NEB, 40 U/µL), 1 μL NAD^+^ (NEB), 1 μL dNTP mix (NEB, 10 mM) and 2 µL 10x NEBuffer 3 (NEB) in a total volume of 20 µL and was incubated for 40 min at 37 °C and 60 min at 45 °C. gDNA was purified using Pronex Magnetic Beads as described previously and eluted in 20 µL.

### HPLC-MS/MS analysis of 8-oxodG in gDNA

40 µg gDNA was used for each sample. Digestion mix was prepared for each sample in a total volume of 50 µL. First, 2.5 mM deferoxamine, 2.5 mM N-*tert*-butyl-α-phenylnitrone, 20 mM Tris-HCl (pH 7.9), 20 mM MgCl2, 100 mM NaCl and LC-MS-grade H2O were mixed. To this mixture, 50 U Benzonase Nuclease (250 U/µL, Sigma), 0.06 U PDE I (0.1 U/µL, US Biologicals), 40 U Antarctic Phosphatase (5 U/µL, NEB), 100 pg [15N5]8oxodG (Cambridge Isotope Laboratories) were added and gently mixed. 50 µL digestion mix was added to 150 µL gDNA and incubated for 4 h at 37 °C while shaking at 300 rpm. Digested samples were loaded on a prewashed 10 kDA MWCO filter unit (VWR) and centrifuged (16500 rcf, 10 min). For total dG quantification 2 µL flow-through was taken. Remaining flow-through was used for 8-oxoG enrichment. Solid phase extraction was performed for 8-oxodG enrichment on a vacuum manifold (Visiprep 24™ DL from Supelco). SPE columns (Strata X 33 M Polymeric RP, 30 mg / 1 ml tubes (8B-S100-TAK) from Phenomenex) were activated twice with 1 mL methanol and twice with 1 mL LC-MS H2O. The remaining flow-through from the previous step was loaded on the activated columns. Columns were washed twice with 1 mL LC-MS-grade H2O. Enriched samples were eluted in 20 % methanol and concentrated to dryness using vacuum centrifugation (miVAC concentrator, DUC-23050-D00). Dried samples were redissolved in 25 μL H2O and sonicated for 10 min (Telsonics Ultrasonics TPC-120).

The liquid chromatography-nanoelectrospray ionization-tandem mass spectrometry (LC-NSI-MS/MS) was performed on a TSQ Quantiva triple quadrupole mass spectrometer (ThermoFisher Scientific, San Jose, CA, United States) coupled to an ACQUITY UPLC M-Class (Waters, Milford, MA, United States) system using nanoelectrospray ionization. The analysis was conducted using a capillary column (150 µm ID, 5.5 cm packing length, 15 µm orifice) created by hand filling a commercially available fused-silica emitter (MSWIL, Aarle-Rixtel, Noord-Brabant, Netherlands) with HSS T3 separation media (Waters, Milford, MA, United States). The mobile phase consisted of 0.1 % (v/v) formic acid in water (A1) and 0.1 % (v/v) formic acid in acetonitrile (B1). A 1 µL injection loop was used and the sample (1 µL) was loaded onto the capillary column with 2 µL/min flow at the initial conditions (95 % A1, 5 % B1) for 2 min and eluted with a linear gradient at a flow rate of 2 µL/min over 3 min to 60 % A1, following by ramping to 99 % B1 within 2 min and holding at this composition for an additional 1 min. The column was then re-equilibrated at the initial conditions for 3 min before the next injection. The nanoelectrospray source was operated in positive ion mode with the voltage set at 2.7 kV. The ion transfer tube temperature was 250 °C and the radio frequency (RF) lens was used as calibrated. The collision gas was argon at 1.5 mTorr with collision energy of 22 eV and the quadrupoles were operated at a resolution of 0.7 Da for both Q1 and Q3. The mass transitions for monitoring the analytes were m/z 268.1 → 152.1 for dG, m/z 284.1 → 168.1 for 8oxodG and m/z 289.1 → 173.1 for [15N5]8oxodG, respectively.

The quantitation of the analytes was done by using the mass spectrometers vendor software package Quan Browser in the software suite Xcalibur based on the peak areas and the constructed calibration curves. Calibration curves were constructed for each analyte during each analysis using a series of standard solutions of analytes. Calibration curves for dG and 8-oxodG were prepared with the concentration of 200 nM, 175 nM, 150 nM, 75 nM, 50 nM, and 25 nM respectively 75 nM, 50 nM, 40 nM, 30 nM, 20 nM, and 15 nM.

### Fluorescence labeling of DNA modifications in gDNA (click-fluoro-quant)

If cells were treated with irofulven, adducts were converted to AP sites by diluting 4 μg gDNA with H_2_O to a final volume of 15 μL and incubating for 18 h at 37°C. The choice of enzymes in the following step depends on the lesion of interest as described in the main text. Enzymes can be replaced by H2O if the analysis of a specific lesion is not desired. To convert DNA modifications to 3’-OH sites, a respective combination of 1 µL FPG (NEB, 8 U/µL), 1 µL ENDOIV (NEB, 10 U/µL) and 1 µL T4 PNK (NEB, 10 U/µL) was mixed with 4 µg gDNA (or a sample from previous blocking/repair step) in 1x NEBuffer 2 (NEB) in a final volume of 20 µL. The reaction mixture was incubated for 1 h at 37°C, immediately followed by adding 3 µL prop-dGTP (Jena Biosciences, 2.5 mM), 3 µL 10x ThermoPol buffer (NEB), 3 µL H2O and Therminator IX (NEB 10 U/µL) to the sample. In case of quantifying AP sites, 3 µL H2O were replaced by 3 µL prop-dATP (Jena Biosciences, 2.5 mM). After 10 min incubation at 60°C, gDNA samples were purified using Pronex® Magnetic Beads as described previously using an elution volume of 30 µL. For CuAAC, the sample volume was condensed to 6 µL using a vacuum concentrator. The reaction was performed using 4 µL AF594 picolyl azide (10 mM stock in DMSO, Jena Biosciences), 4 µL DMSO, 2 µL premixed CuSO4:THPTA (THPTA from Lumiprobe; 20 mM Cu^2+^ and 200 mM THPTA) and 2 µL sodium phosphate buffer (1 M, pH 7.0). The reaction was started by adding 2 µL sodium ascorbate (400 mM in H2O, freshly prepared) and incubated for 60 min at 37°C. The reaction was purified using the Monarch Genomic DNA Purification Kit and eluted in 100 µL Buffer EB. gDNA concentration was measured using the Quantus™ Fluorometer with QuantiFluor® ONE dsDNA dye. For fluorescence analysis, identical amounts of each gDNA sample were mixed with PBS for a final volume of 605 µL. The samples were split in 3x 200 µL and pipetted in a non-clear 96 black well plate. For blank measurement, 3x 200 µL PBS were added to the plate. Fluorescence was recorded with an Infinite Pro M200 Plate Reader (Tecan) using the following settings: λEx = 580 nm, λEm = 610 nm, gain = optimal, 25 flashes, 20 µs integration time, 25°C. The same gain value was used for replicates of the same experiment and usually set to 255. Data were analyzed by subtracting the median blank fluorescence from all sample fluorescence values. They were further normalized to the DNA amount by dividing each value by the amount of gDNA in the sample in ng. The resulting values were rescaled by min-max normalization using the formula 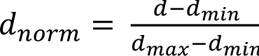, where *d* is a datapoint, *d_min_* the lowest value and *d_max_* the highest value in a dataset.

### Plotting and statistics

For all plots concerning click-fluoro-quant and related statistical analyses, we used Jupyter notebooks employing the modules numpy version 1.22.4, scipy version 1.8.1, pandas version 1.4.2, matplotlib version 3.5.2 and seaborn version 0.11.2 in Python version 3.10.4 (main, May 30 2022, 08:01:42) [GCC 8.2.0]. The code: https://gitlab.ethz.ch/eth_toxlab/click-code-seq/, folder: click-fluoro-quant. See below the details of click-code-seq data analysis.

### Click-code-seq library preparation

To minimize artefactual damage, samples were inverted or flicked, instead of pipette mixing, until the CuAAC step. In case of endogenous-DNA-oxidation mapping, gDNA was eluted in H2O supplemented with 500 µM 8-hydroxyquinoline (HQ; Sigma) after extraction and purification. Undesired background modifications were blocked by adding 1 µL ENDOIV (NEB, 10 U/µL) and 500 µM 8-HQ to 5 µg of gDNA, freshly extracted from HAP1 pellets, in 1x ThermoPol Buffer (NEB) in a final volume of 20 µL, and incubating the mixture for 20 min at 37°C. 2.5 µL ddNTP mix (Jena Biosciences, 2.5 mM), 0.5 µL 10x ThermoPol Buffer, 1.8 µL of 8-HQ (500 µM) and 0.2 µL Therminator IX (NEB, 10 U/µL) were added, followed by a 10 min incubation at 60°C. 25 µL 8-HQ (500 µM) and 80 µL Pronex® beads were added for purification with an elution volume of 25 µL. To label oxidative modifications, 1 µL FPG (NEB, 8 U/µL), 1 µL ENDOIV (NEB, 10 U/µL) and 3 µL 10x NEBuffer 2 (NEB) were added for a final volume of 30 µL; the reaction mixture was incubated for 1 h at 37°C, immediately followed by adding 3.5 µL prop-dGTP (Jena Biosciences, 2.5 mM), 0.5 µL 10x NEBuffer 2 (NEB), 0.8 µL 8-HQ (500 µM) and 0.2 µL Therminator IX (NEB 10 U/µL) for 10-min incubation at 60°C.

In case of mapping AP sites, 5 µg of freshly extracted U2OS gDNA was diluted with water to a final volume of 20 µL and incubated at 37°C for 18 h; then, to dephosphorylate background strand breaks, 1 µL T4 polynucleotide kinase (T4 PNK) (10 U/µL, NEB) was added to the gDNA in 1x NEBuffer 2 (NEB) in a final volume of 25 µL and incubated for 30-min at 37°C. 3 µL ddNTP mix (Jena Biosciences, 2.5 mM), 0.5 µL 10x NEBuffer 2 (NEB), 1.3 µL water and 0.2 µL Therminator IX (NEB, 10 U/µL) were added, followed by a 10 min incubation at 60°C. 20 µL water and 80 µL Pronex® beads were added for purification with an elution volume of 25 µL. To label AP sites, 1 µL Endonuclease IV (ENDOIV) (10 U/µL, NEB) and 3 µL 10x ThermoPol buffer (NEB) were added for final volume of 30 µL; the reaction mixture was incubated for 30 min at 37°C, immediately followed by adding 4 µL prop-dGTP (2.5 mM in H2O, Jena Biosciences), 5 µL prop-dATP (2 mM in H2O, Jena Biosciences), 1 µL 10x ThermoPol buffer (NEB), 0.8 µL H2O and 0.2 µL Therminator IX (10 U/µL, NEB) for 10-min incubation at 60°C.

Steps for mapping either AP sites or oxidative damage are identical from this point onwards. gDNA was purified using Pronex® Magnetic Beads (Promega) with a sample:bead ratio of 50:80 and using an elution volume of 50 µL with Buffer EB. Sonication of gDNA was performed using a Q800R2 sonicator (QSonica) with the following settings: 4°C, 20% amplitude, pulse 15 s on, 5 s off, total 5 min. The resulting DNA fragments were analyzed on a Tapestation 2200 using a High Sensitivity D100 ScreenTape (Agilent Technologies). DNA was purified using Pronex® beads with a sample:beads ratio of 1:2 and 50 µL elution volume. Samples were quantified using a Quantus™ Fluorometer. For adapter ligation, NEBNext® Ultra™ II DNA Library Prep Kit for Illumina® (NEB) was utilized using 1 µg gDNA and the pre-annealed NEB adapter (NEB_P7, NEB_P5, 15 µM; Supplementary Table 1). The manufacturer’s instruction was followed without performing the USER enzyme step. The sample was purified using Pronex® beads in sample:beads ratio 1:2 and elution in 50 µL Buffer EB. Successful adapter ligation was checked on an Agilent TapeStation with a High Sensitivity D100 ScreenTape. The sample volume was then condensed to 6 µL using a vacuum concentrator. To denature gDNA, the sample was heated to 95°C for 2 min and rapidly cooled on ice. CuAAC reaction was performed using 2 µL MoDIS (100 µM equimolar mixture of MoDIS_1 and MoDIS_2; Supplementary Table 1), 4 µL DMSO, 2 µL premixed CuSO4:THPTA (THPTA from Lumiprobe; 20 mM Cu^2+^ and 200 mM THPTA) and 2 µL sodium phosphate buffer (1 M, pH 7.0). The reaction was started by adding 2 µL sodium ascorbate (400 mM in H2O, freshly prepared) and incubated for 60 min at 37°C. Purification was done using Pronex® beads in a sample:beads ratio 1:2 and 30 µL elution volume. Biotin enrichment of sample DNA was performed using Dynabeads™ MyOne™ Streptavidin C1 (ThermoFisher). First, B&W buffer was prepared according to the manufacturer’s protocol including 0.05 % Tween™ 20. Then, 15 µL Dynabeads™ were transferred to a fresh tube, washed three times with 200 µL 1x B&W buffer and resuspended in 30 µL 2x B&W buffer. The DNA sample was denatured by heating to 95°C for 2 min and rapid cooling on ice. Then, 30 µL sample was mixed with 30 µL of the prepared Dynabeads™ and rotated for 15 min at RT. Afterwards, the Dynabeads™ were washed 3x with 1x B&W buffer by resuspending the beads, 3 min rotation and 1 min pelleting on a magnetic rack. A final wash was performed with Buffer EB and Dynabeads™ were resuspended in 20 µL Buffer EB. To amplify the biotin-enriched material, a PCR was performed by adding 5 µL 10x ThermoPol buffer, 5 µL dNTP mix (NEB, 2 mM), 1 µL MgSO4 (100 mM), 1 µL ex_UMI (10 µM; Supplementary Table 1), 1 µL Pr_P7 (10 µM; Supplementary Table 1) and 0.5 µL Vent (exo-) DNA Polymerase (NEB 2 U/µL). The PCR program comprised an initial denaturation step (95°C, 120 s), followed by 5 cycles of denaturation (95°C, 20 s), annealing (59°C, 20 s) and elongation (72°C, 60 s) and a final elongation step (72°C, 300 s). The sample was purified with Pronex® beads in a sample:beads ratio 1:2 and an elution volume of 30 µL. Final library amplification and indexing was performed by adding 25 µL NEBNext® Ultra™ II Q5® Master Mix (NEB), 2.5 µL i5_Index (10 µM; Supplementary Table 1) and 2.5 µL i7_Index (10 µM; Supplementary Table 1) to 20 µL sample. The PCR program comprised an initial denaturation step (98°C, 30 s), 10 cycles of denaturation (98°C, 10 s), annealing (68°C, 30 s) and elongation (72°C, 20 s), and a final elongation step (72°C, 120 s). The same PCR setup can be used for a precedent qPCR by adding a qPCR dye as Evagreen (Biotium) to test for successful sample amplification. The PCR sample was purified with the Pronex® beads dual size selection protocol using an initial sample:beads ratio of 1:1, followed by a sample:beads ratio of 0.5:1 to exclude fragments longer than 1000 bp and shorter than 250 bp. The final sample concentration was adjusted to a concentration of 4 ng/µL and sequenced on an Illumina NovaSeq 6000.

### Click-code-seq data analysis

#### Sequencing read processing

After demultiplexing of sequencing data, each sample was represented by a fastq.gz file containing 101-nucleotide-long genomic reads. The quality of the raw sequencing data was checked via FastQC version 0.11.9. Low-quality reads and adapter-containing reads were removed via trimmomatic version 0.38 with the following parameters: SE ILLUMINACLIP:Trimmomatic-0.39/adapters/TruSeq3-SE.fa:2:30:10 LEADING:3 TRAILING:3 SLIDINGWINDOW:4:15 MINLEN:101. We retained only those reads that contained a validation code (VC). The first 17 nucleotides corresponding to VC and randomized index code (RIC) were clipped from the read sequences and appended to the read names via the tool extract of umi_tools/1.1.2 toolkit. The reads were mapped to human reference genome GRCh38 via bowtie2 version 2.3.5.1, using pre-built bowtie2 index from https://genome-idx.s3.amazonaws.com/bt/GRCh38_noalt_as.zip, and applying otherwise standard settings. Read duplicates were removed by the tool *dedup* of umi_tools version 1.1.2 toolkit, grouping reads with the same code sequence stored in the read name (*method=unique*). Samtools version 1.12 were employed to sort, index and generate statistics of bam files. bedtools2 version 2.29.2 were used to retrieve the coordinates of mapped and deduplicated reads and extract the sequence context of the damaged nucleotides from the reference genome. Each read represented one unit of DNA-modification signal, which we positioned at the nucleotide of the 5’ end of the read. Since a read was the reverse complement of the DNA fragment captured in the method employing MoDIS, the strand of the nucleotide bearing the signal was changed to the opposite one. Supplementary Fig. 2g describes the evolution of read counts throughout the preprocessing steps. We implemented the described DNA-modification-signal positioning via custom scripts in Python version 3.7.4 (default, Oct 3 2019, 08:07:56) [GCC 4.8.5 20150623 (Red Hat 4.8.5-28)] with the modules numpy version 1.21.5, pandas version 0.25.1 and biopython version 1.79.

#### Software for mapped DNA-modification data analysis

The downstream analysis of DNA-modification data and their visualization were performed in Jupyter notebooks employing the modules numpy version 1.26.4, scipy version 1.13.1, pandas version 2.2.2, biopython version 1.83, matplotlib version 3.9.0, seaborn version 0.13.2 and pyfaidx version 0.8.1.1 in Python version 3.11.6 (main, Jun 7 2024, 07:09:59) [GCC 13.2.0]; in Jupyter notebooks employing the modules numpy version 1.22.4, scipy version 1.8.1, pandas version 1.4.2, biopython version 1.79, matplotlib version 3.5.2, seaborn version 0.11.2, logomaker version 0.8 and pyfaidx version 0.6.4 in Python version 3.10.4 (main, May 30 2022, 08:01:42) [GCC 8.2.0]; in Jupyter notebooks employing the modules numpy version 1.19.2 and pandas version 1.1.3 in Python version 3.8.5 (default, Oct 6 2020, 10:04:29) [GCC 6.3.0]; in custom scripts in Python version 3.7.4 (default, Oct 3 2019, 08:07:56) [GCC 4.8.5 20150623 (Red Hat 4.8.5-28)] with the modules numpy version 1.21.5, pandas version 0.25.1 and biopython version 1.79. The custom code for analyzing click-code-seq data and plotting figures is available at https://gitlab.ethz.ch/eth_toxlab/click-code-seq.

#### External datasets

Human reference genome, GRCh38 [https://genome-idx.s3.amazonaws.com/bt/GRCh38_noalt_as.zip] (bowtie2 pre-built index). For the analysis related to gene expression and chromatin accessibility, we used the data from GRCh38 chr1-22 and chrX. Single base substitution signatures: COSMIC v3.4 [https://cancer.sanger.ac.uk/signatures/documents/2124/COSMIC_v3.4_SBS_GRCh38.txt]. Illudin S mutational signature: study ^56^ [https://ars.els-cdn.com/content/image/1-s2.0-S1568786422001665-mmc1.xlsx], Supplementary material, Tab_S4, ILS Clones, % of SBS (Count no background), ILS Sig. Centromeres and gaps: UCSC Table Browser for GRCh38. Chromatin accessibility and histone marks: ChIP-Atlas [https://chip-atlas.org/peak_browser], significance threshold 50, downloaded on 13.02.2024, with the following accession numbers for U2OS: H3K4me1 - GSM1901957, GSM4861712; H3K9me3 - GSM4079837, GSM3147771, GSM788078; H3K4me3 - GSM2341637, GSM3147770; H3K36me3 - GSM788076; H3K27ac - GSM2341636, GSM6836721, GSM4133285, GSM4133286, GSM5018581, GSM5018582, GSM6836725; DNase hypersensitive sites - GSM4221655; for HAP1: H3K4me1 - GSM3579008; H3K9me3 - GSM3579026; H3K4me3 - GSM5570286, GSM2978165, GSM3901520, GSM2978166, GSM3901519; H3K36me3 - GSM5570287; H3K27me3 - GSM5570291, GSM2978164, GSM3901518, GSM3901517, GSM2978163, GSM5770668, GSM5770667; H3K27ac - GSM3901521, GSM3901522; DNase hypersensitive sites - GSM5214993, GSM5214992, GSM2400413, GSM2400414. Transcript coordinates: GENCODE/V41/knownGene, obtained from UCSC Table Browser. Canonical transcripts of genes: GENCODE/V41/knownCanonical, obtained from UCSC Table Browser. Protein-coding genes: GENCODE/V41/knownToNextProt, obtained from UCSC Table Browser. Genes were represented by canonical transcripts between transcription start site (TSS) and transcription end site (TES). Gene expression: CCLE_expression_full from DepMap Public 22Q2 (https://depmap.org/portal/download/all/), the cell-line accession numbers: ACH-002475 (HAP1) and ACH-000364 (U2OS). Mitochondrial gene and D-loop-region annotation: NCBI Reference Sequence NC_012920.1 [https://www.ncbi.nlm.nih.gov/nuccore/251831106].

#### Correlation between chromatin state and DNA modifications

For every bin with its lower and upper boundaries *b* = [*b_lo_*, *b_up_*] and ChIP-Seq mapping study *z* with chromatin-mark peaks defined by the lower and upper boundaries *p*(*z*) = [*b_lo_*, *b_up_*], we calculated the peak coverage of the bin 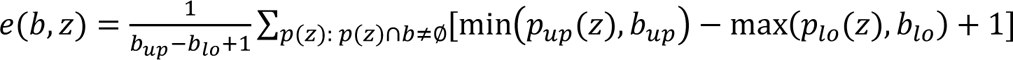. We excluded bins that do not contain the target nucleotide (G or A) according to the reference genome or overlap with centromeric regions or heterochromatin and short-arm gaps of the reference genome. For each type of DNA modification *T*, namely, guanine oxidation, irofulven-induced AP sites at adenines or background AP sites at guanines, we averaged the binned DNA-modification data 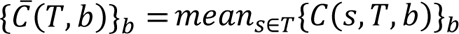, where *s* is a replicate experiment and *C*(*s, b*) is the reference-genome- and sequencing-depth-normalized DNA-modification level per bin (see the formula below in the gene-related analysis). For each ChIP-Seq mapping study *z* and DNA-modification type *T*, we then calculated the coefficient of Spearman correlation *ρ*(*z*, *T*) between the binned chromatin-mark data {*e*(*b*, *z*)}*_b_* and the averaged binned DNA-modification data 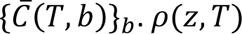 are presented as markers in Fig.3, with the bar being the median across the mapping studies *z*.

#### DNA-modification level normalization, gene-related analysis

DNA-modification levels in genomic features of interest were corrected by respective counts of target nucleotides (G or A) in the reference genome. Additionally, the genome-scale data required normalization to correct for sequencing depth that varied among samples (Supplementary Fig. 2g). We therefore computed sample-specific normalization factors that reflect the level of DNA modification *T* (oxidized guanines, adenosine-derived AP sites or guanosine-derived AP sites) in unexpressed genes: *M*(*s, T*) = median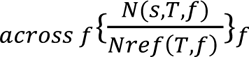, where features *f* here are the transcribed (antisense) strands of unexpressed protein-coding genes, *N*(*s*, *T*, *f*) is the count of mapped DNA modifications *T* per a concrete feature *f* in the sample *s*, and *Nref*(*T, f*) is the feature’s count (kilobases) of target nucleotides for *T* in the reference genome. DNA-modification levels *C*(*s, T, f*) were related to these sample-specific normalization factors in the following way: 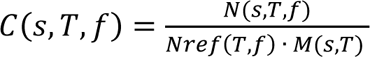 [arb. unit], where a feature *f* is a dinucleotide (Fig. 2d), a chromosome bin with both strands considered together (Fig. 3, Supplementary Fig. 2h, 3c,f,i,k-m), a gene’s transcribed strand or a gene’s non-transcribed strand (Fig. 4a,b,e,f,i,j, Supplementary Fig. 5a,b,e,f), strands of the TSS-adjacent region (Fig. 4m-p, Supplementary Fig. 8). DNA-modification strand bias was calculated via *C*(*s, T, f_NTS_*) − *C*(*s, T, f_TS_*), where *f_NTS_* and *f_TS_* are respectively non-transcribed and transcribed strands of the same feature (Fig. 4d,h,l, Supplementary Fig. 5d,h). For metaprofiles (Fig. 4c,g,k, Supplementary Fig. 4b-d, Supplementary Fig. 5c,g), we presented 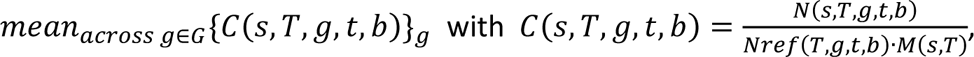 where *N*(*s*, *T*, *g*, *t*, *b*) is the count of mapped DNA modifications *T* per bin *b* in sample *s*, gene *g* and strand *t*, *Nref*(*T, g, t, b*) is the bin’s count (kilobases) of target nucleotides for *T* in reference genome in gene *g* and strand *t*, *M*(*s, T*) is the normalization factor defined above, and *G* is the set of 30% most expressed or unexpressed protein-coding genes. In trinucleotide-specific analysis (Supplementary Figs. 4a, 6 and 7), DNA-modification levels *C*(*s, T, g, t*) were related to the maximal median value within a sample (the maximum of the median values for each sample shown in the plots is set to 1): *C*(*s, T, g, t*)/*max_across(G,t)_*{*median_acrossgεG_*{*C*(*s, T, g, t*)*_g_*}*_G,t_* where *g* is a gene and *t* is a strand, such that occurrences of only a certain trinucleotide within this gene and strand are considered, *G* is a gene-expression tier, (*G, t*) is a combination of gene expression tier and strand. In mDNA analysis (Fig. 5, Supplementary Fig. 9d), DNA-modification level was calculated as follows: *C*(*s*, *T*, *t*, *b*) = 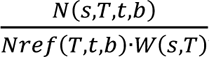, where *N*(*s*, *T*, *t*, *b*) is the count of mapped DNA modifications *T* per bin *b* in sample *s* and strand *t*, *Nref*(*T*, *t*, *b*) is the bin *b*’s count (bases) of target nucleotides for *T* in strand *t* according to the reference genome, *W*(*s*, *T*) is the total count of mapped DNA modifications *T* in mDNA in sample *s*.

In HAP1, we considered 16,740 protein-coding genes, including unexpressed ones: 3,428; ≤ 10% expression tier: 1,404; ≤ 20%: 1,259, ≤ 30%: 1,332, ≤ 40%: 1,330, ≤ 50%: 1,333, ≤ 60%: 1,329, ≤ 70%: 1,331; ≤ 80%: 1,331; ≤ 90%: 1,331; ≤ 100%: 1,332. Number of data points beyond the maximal Y-axis value in Fig. 3a, 2,202 (TS), 2,228 (NTS); in Fig. 3d, 48; in Fig. 4m, 506 (TS), 493 (NTS). In U2OS, we considered 16,740 protein-coding genes, including unexpressed ones: 1,989; ≤ 10% expression tier: 1,544; ≤ 20%: 1,415, ≤ 30%: 1,467, ≤ 40%: 1,478, ≤ 50%: 1,474, ≤ 60%: 1,474, ≤ 70%: 1,474; ≤ 80%: 1,475; ≤ 90%: 1,475; ≤ 100%: 1475. Number of data points beyond the maximal Y-axis value in Fig. 4e, 396 (TS), 526 (NTS); Supplementary Fig. 5e, 2,624 (TS), 3,112 (NTS); Fig. 4i, 1,116 (TS), 1,348 (NTS); Supplementary Fig. 5a, 502 (TS), 528 (NTS); Fig. 4h, 40; Supplementary Fig. 5h, 2,485; Fig. 4l, 251; Supplementary Fig. 5d, 79; Fig. 4o, 995 (TS), 959 (NTS); Supplementary Fig. 8a, 3,239 (TS), 3,277 (NTS); Supplementary Fig. 8c, 1,319 (TS), 1,346 (NTS); Supplementary Fig. 8e, 2,361 (TS), 2,399 (NTS).

## Data availability

The raw sequencing data and processed sequencing data (tsv-files with called DNA-modification sites) generated in this study are deposited in the NCBI Gene Expression Omnibus (GEO), GEO series accession code: GSE272366 [https://www.ncbi.nlm.nih.gov/geo/query/acc.cgi?acc=GSE272366].

## Code availability

The code of the data analysis required to generate the figures is available at GitLab [https://gitlab.ethz.ch/eth_toxlab/click-code-seq].

## Acknowledgements

Genome-scale data produced and analyzed in this paper were generated using the resources of the ETH Zürich Genetic Diversity Centre (GDC), the Functional Genomics Center Zürich (FGCZ), and the ETH Zürich Euler cluster. The authors thank Emma Dillier and Chantal Balmer from ETH Zürich Laboratory of Toxicology for experimental support, Andrew Gardner, Karl von Laer and Claudia Lehmann from New England Biolabs for advice on enzyme applications, Thomas Nicholls from Newcastle University for providing valuable insights into mitochondrial genetics. Fig. 1a was made with the help of biorender.com. This work was supported by the Swiss National Science Foundation (grants #185020 and #186332 to SJS).

## Author contributions

VT, NJLP, HG and SJS designed the study. NJLP performed experiments with oligonucleotide DNA and optimized conditions for experiments with genomic DNA. NKS performed experiments on genomic DNA. SH and SS performed LC-MS/MS analysis. VT designed the bioinformatics pipeline and performed analysis of sequencing data. SJS, VT and ARP contributed to the evaluation of data, experimental planning, and supervision of research. SJS acquired funding. VT, NJLP and SJS wrote the manuscript.

## Supplementary information

PDF contains Supplementary Figures 1–9, Supplementary Table 1.

