## Supplementary Information for "Click-chemistry-aided quantitation and sequencing of oxidized guanines and apurinic sites uncovers their transcription-linked strand bias in human cells"

**This file includes:**

- Supplementary Figures 1–9, pages 2–13
- Supplementary Table 1, page 14
- Supplementary Note 1, page 15

### Supplementary Figures

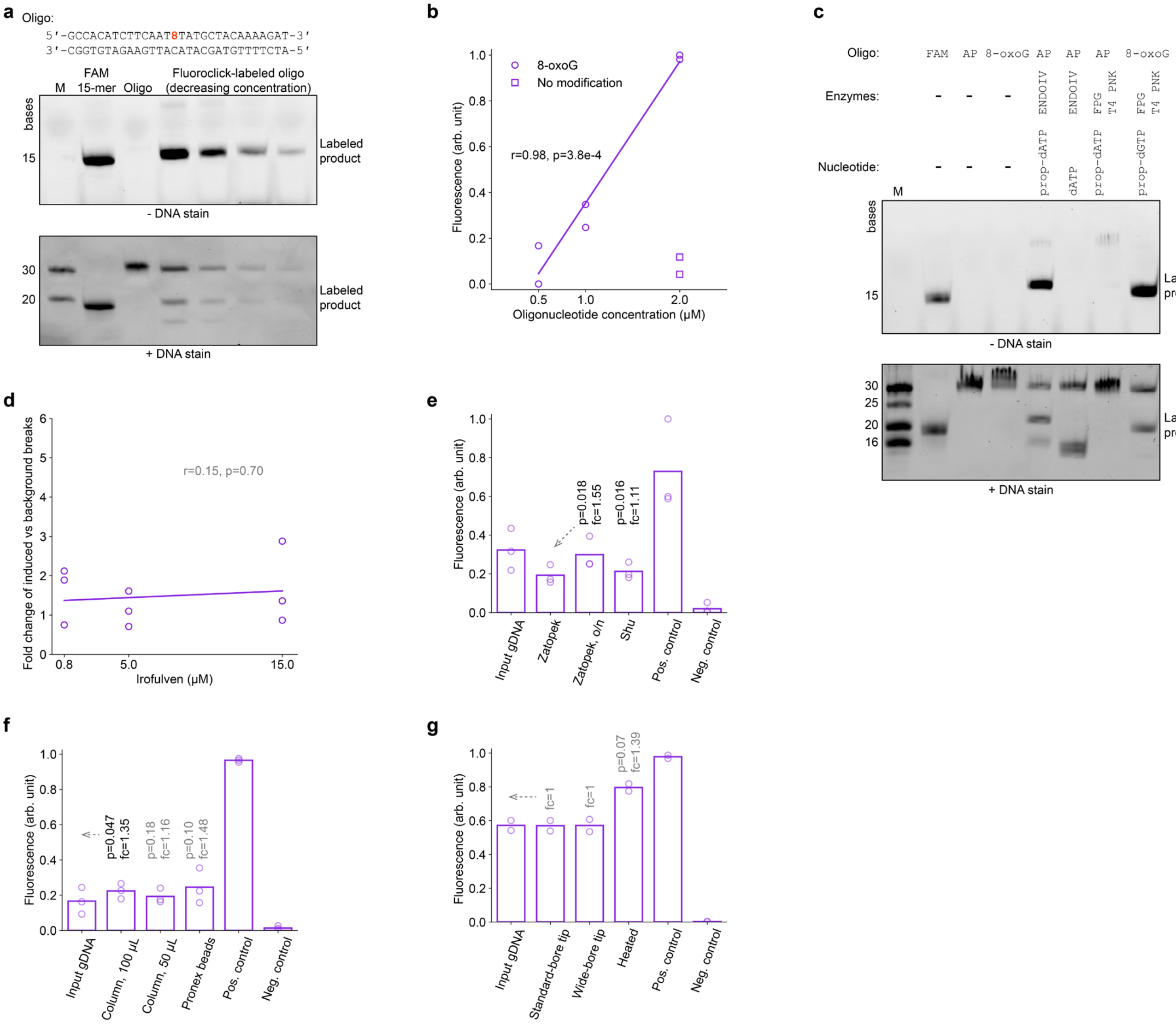


**Supplementary Fig. 1 | Click-fluoro-quant to assess DNA-modification levels in oligonucleotides and gDNA. a** Products of click-fluoro-quant labeling of 8-oxoG site at position 15 in the double-stranded 30-mer DNA oligonucleotide in a PAGE gel. 8, 8-oxoG. Labeled product: AF594-labeled 15-mer fragment. The unmodified 30-mer complimentary strand and the unlabeled 15-mer fragment originating from 8-oxoG excision are visible in DNA staining. AF594 conjugation causes a higher retention time resulting in a band corresponding to a strand of 20 bases. **b** Fluorescence intensity of click-fluoro-quant-labeled oligonucleotides containing 8-oxoG (**a**) and lacking this DNA modification. n=2 independent experiments. **c** Click-fluoro-quant labeling of a 30-mer oligonucleotide containing an AP-site-mimicking tetrahydrofuran at position 17 or 8-oxoG at position 15 (**a**) in a PAGE gel. **d** Click-fluoro-quant of irofulven-induced 3'-OH breaks in U2OS cells after 4-h exposure to the drug. Extracted gDNA was incubated overnight at 37 °C (to induce depurination of irofulven adducts, Fig. 1f) and directly labeled in fluoroclick without enzyme treatment. Markers: biological replicates (n=3), each of which included three drug concentrations and a vehicle control by which the plotted values are normalized. **e** Click-fluoro-quant comparison of gDNA repair efficiencies of three protocols: according to Zatopek *et al.* (also used in Fig. 1g), Zatopek *et al.* with overnight (o/n) enzymatic repair and Shu *et al.* **f-g** Impact of sample handling on DNA-modification levels in HAP1 gDNA. **f** Click-fluoro-quant after gDNA purification with silica columns using two different elution volumes or magnetic beads. **g** Click-fluoro-quant after pipetting gDNA 20 times with a standard-bore or a wide-bore tip, or after incubating gDNA at 85 °C for 20 min (Heated). In panels **a,c**, the gel was imaged in the fluorophore and DNA-stain channels before and after applying GelRed® DNA stain, respectively; M, marker; FAM, FAM-labeled 15-mer oligonucleotide.

**b,d**: r and p: Pearson's correlation coefficient with respective p-value. In panels **e-g**, bar: mean of n=2 (**f-g**) or n=3 (**e**) biological replicates (markers); p: p-value of the one-tailed ($H_A$: greater) paired t-test between the samples under the value and indicated by the arrow; fc: fold change of DNA-modification level relative to the arrow-indicated condition; Pos. control: gDNA incubation with FPG and ENDOIV; Neg. control: dGTP incorporation instead of prop-dGTP.

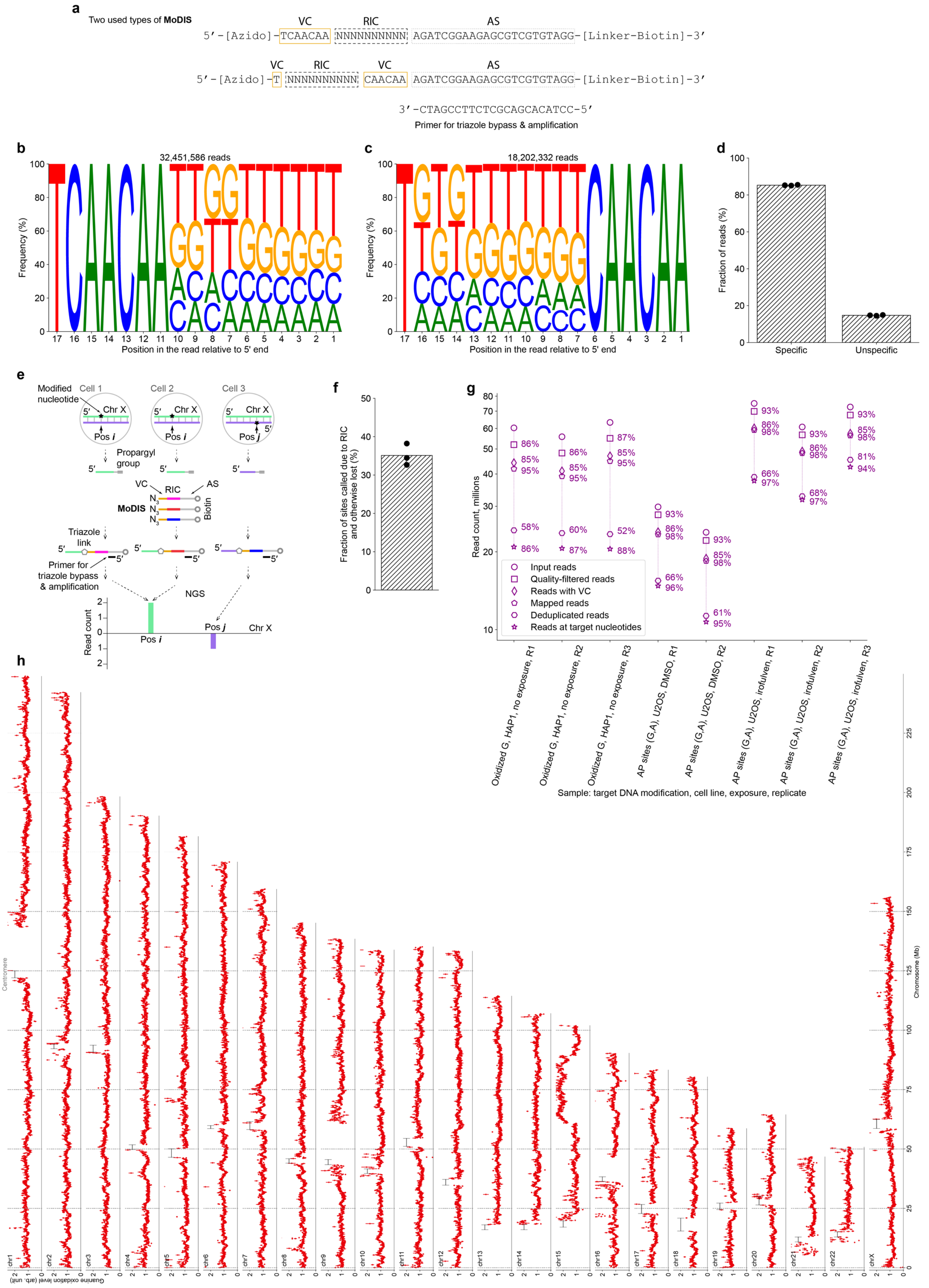
a
Two used types of MoDIS
VC
RIC
AS
5′-[Azido]-TCAACAA NNNNNNNNNN AGATCGGAAGAGCGTCGTGTAGG-[Linker-Biotin]-3′
VC
RIC
VC
AS
5′-[Azido]-T NNNNNNNNNN CAACAA AGATCGGAAGAGCGTCGTGTAGG-[Linker-Biotin]-3′
3′-CTAGCCTTCTCGCAGCACATCC-5′
Primer for triazole bypass & amplification
b
32,451,586 reads
Frequency (%)
Position in the read relative to 5′ end
c
18,202,332 reads
Frequency (%)
Position in the read relative to 5′ end
d
Fraction of reads (%)
Specific
Unspecific
e
Modified nucleotide
Cell 1
Cell 2
Cell 3
Chr X
Pos i
Pos j
Propargyl group
VC
RIC
AS
MoDIS
Biotin
Triazole link
Primer for triazole bypass & amplification
NGS
Read count
f
Fraction of sites called due to RIC and otherwise lost (%)
g
Read count, millions
Input reads
Quality-filtered reads
Reads with VC
Mapped reads
Deduplicated reads
Reads at target nucleotides
Oxidized G, HAP1, no exposure, R1
Oxidized G, HAP1, no exposure, R2
Oxidized G, HAP1, no exposure, R3
AP sites (G,A), U2OS, DMSO, R1
AP sites (G,A), U2OS, DMSO, R2
AP sites (G,A), U2OS, irofulven, R1
AP sites (G,A), U2OS, irofulven, R2
AP sites (G,A), U2OS, irofulven, R3
Sample: target DNA modification, cell line, exposure, replicate
h
Centromere
Chromosome (Mb)
Guanine oxidation level (arb. unit)
chr1
chr2
chr3
chr4
chr5
chr6
chr7
chr8
chr9
chr10
chr11
chr12
chr13
chr14
chr15
chr16
chr17
chr18
chr19
chr20
chr21
chr22
chrX

**<<< Supplementary Fig. 2 | The upgraded design of click-code-seq for single-nucleotide-resolution DNA-modification sequencing.** Modified DNA Identifier Sequence (MoDIS) helps filtering out unspecific reads and retaining identical reads originating from different cells/genomes. **a** Two used types of MoDIS. VC, validation code; RIC, randomized index code; AS, annealing site sequence. **b-c** VC (conserved positions) and RIC (variable positions) in the reads originating from MoDIS. The read sequence was complemented. Data: one replicate of endogenous-guanine-oxidation mapping is shown. **d** The fractions of specific, *i.e.*, MoDIS-containing reads (with VC, see **a-c**), and unspecific reads lacking VC. The reads in **b-c** correspond to one circular marker in the group of specific reads. Due to using MoDIS, the unspecific, *i.e.*, artefactual reads (<15%), can be identified and removed. **e** Schematic explanation of how RIC helps retain identical reads originating from different cells/genomes. **f** The usage of RIC in MoDIS elevates the number of identified oxidation sites by around 35%. The fraction relates the number of specific (*i.e.*, VC-containing) deduplicated reads with RIC sequences and the number of reads after computational removal of RIC sequences and deduplication. In panels **d,f**, n=3 biological replicates (circular markers) and their means (bars) of endogenous-oxidized-guanine mapping are shown. **g** Read counts throughout the steps of sequencing-data preprocessing in all samples used in this work. Percentage: fraction of reads remaining after the respective processing step. VC: validation code (**a-c**). Mapping efficiency is 95-98% across the samples. **h** Genome-wide distribution of endogenous guanine oxidation levels at 100-Kb resolution in HAP1 cells; shown: mean ± s.d. (marker and error bar) across n=3 biological replicates. Max Y-axis value: 99.5 percentile of the mean values.

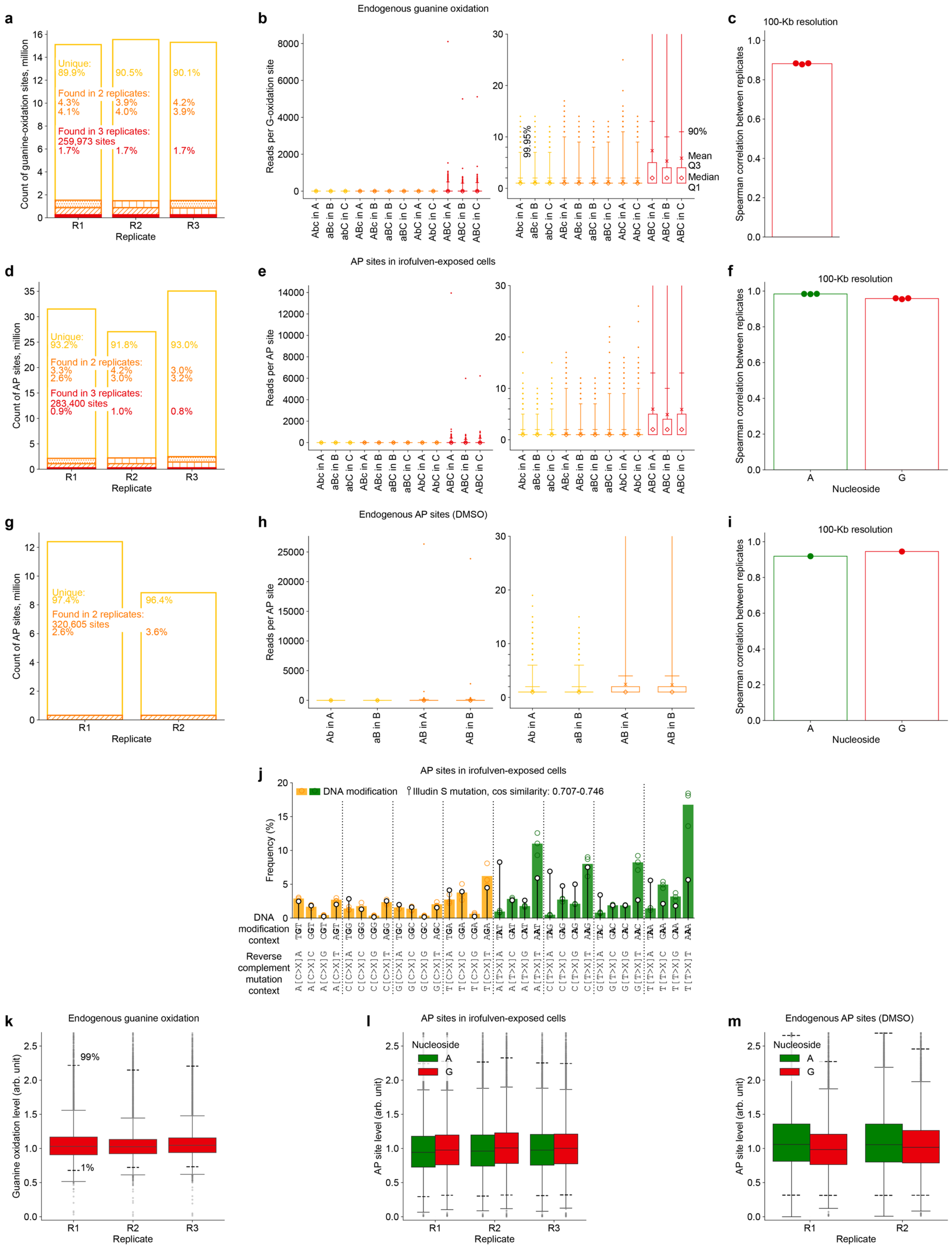
a
Count of guanine-oxidation sites, million
Unique:
89.9%
90.5%
90.1%
Found in 2 replicates:
4.3%
4.1%
3.9%
4.0%
4.2%
3.9%
Found in 3 replicates:
259,973 sites
1.7%
1.7%
1.7%
R1
R2
R3
Replicate
b
Endogenous guanine oxidation
Reads per G-oxidation site
99.95%
90%
Mean
Q3
Median
Q1
Abc in A
aBc in B
abC in C
ABc in A
ABc in B
aBC in B
aBC in C
AbC in A
AbC in C
ABC in A
ABC in B
ABC in C
c
100-Kb resolution
Spearman correlation between replicates
d
Count of AP sites, million
Unique:
93.2%
91.8%
93.0%
Found in 2 replicates:
3.3%
2.6%
4.2%
3.0%
3.0%
3.2%
Found in 3 replicates:
283,400 sites
0.9%
1.0%
0.8%
R1
R2
R3
Replicate
e
AP sites in irofulven-exposed cells
Reads per AP site
f
100-Kb resolution
Spearman correlation between replicates
A
G
Nucleoside
g
Count of AP sites, million
Unique:
97.4%
96.4%
Found in 2 replicates:
320,605 sites
2.6%
3.6%
R1
R2
Replicate
h
Endogenous AP sites (DMSO)
Reads per AP site
Ab in A
aB in B
AB in A
AB in B
i
100-Kb resolution
Spearman correlation between replicates
A
G
Nucleoside
j
AP sites in irofulven-exposed cells
DNA modification
Illudin S mutation, cos similarity: 0.707-0.746
Frequency (%)
DNA modification context
Reverse complement mutation context
k
Endogenous guanine oxidation
Guanine oxidation level (arb. unit)
99%
1%
R1
R2
R3
Replicate
l
AP sites in irofulven-exposed cells
Nucleoside
A
G
AP site level (arb. unit)
R1
R2
R3
Replicate
m
Endogenous AP sites (DMSO)
Nucleoside
A
G
AP site level (arb. unit)
R1
R2
Replicate

**<<< Supplementary Fig. 3 | Reproducibility of DNA-modification signals at the single-nucleotide level and at 100-Kbp resolution; irofulven-induced AP-site profile versus illudin S mutational signature. a,d,g** Counts of single-nucleotide sites of endogenous guanine oxidation in HAP1 cells (**a**), depurination in irofulven-exposed U2OS cells (**d**) and depurination in vehicle-(DMSO)-exposed U2OS cells (**g**) after categorizing the sites into replicate-specific and reproducible across replicate experiments. **b,e,h** The distribution of the number of unique reads (with individual RICs) supporting the DNA modification sites across the replicate experiments and the categories of modification-site reproducibility. A, replicate R1; B, replicate R2; C, replicate R3; Abc, the DNA modification sites specific to replicate A (R1); ABc, the DNA modification sites found in replicates A (R1) and B (R2); ABC, the DNA modification sites found in three replicates. The box plot elements are explained in **b**. **c,f,i** Pair-wise correlations (markers) among n=3 (**c,f**) or n=2 (**i**) replicates for the mapping of endogenous guanine oxidation in HAP1 cells (**c**), AP sites in irofulven-exposed U2OS cells (**f**) and AP sites in vehicle-(DMSO)-exposed U2OS cells (**i**) at 100-Kbp resolution; A: adenosine-derived, G: guanosine-derived AP sites. Bars: respective mean pair-wise correlations. **j** The alignment between Illudin S mutational signature and the distribution of AP sites regarding trinucleotide context in U2OS cells exposed to irofulven (n=3). The plot is built analogously to Fig. 3g. **k-m** The 100-Kbp-bin distributions of endogenous guanine oxidation in HAP1 cells (**k**, the same data as in Supplementary Fig. 2h), AP sites in irofulven-exposed U2OS cells (**l**) and AP sites in vehicle-(DMSO)-exposed U2OS cells (**m**) in individual replicates; A: adenosine-derived, G: guanosine-derived AP sites. Dashed lines: $1^{st}$ and $99^{th}$ percentiles. Boxes are interquartile ranges, internal horizontal lines are medians, whiskers extend to the furthest datapoint within 1.5x interquartile range, datapoints beyond are shown as small markers.

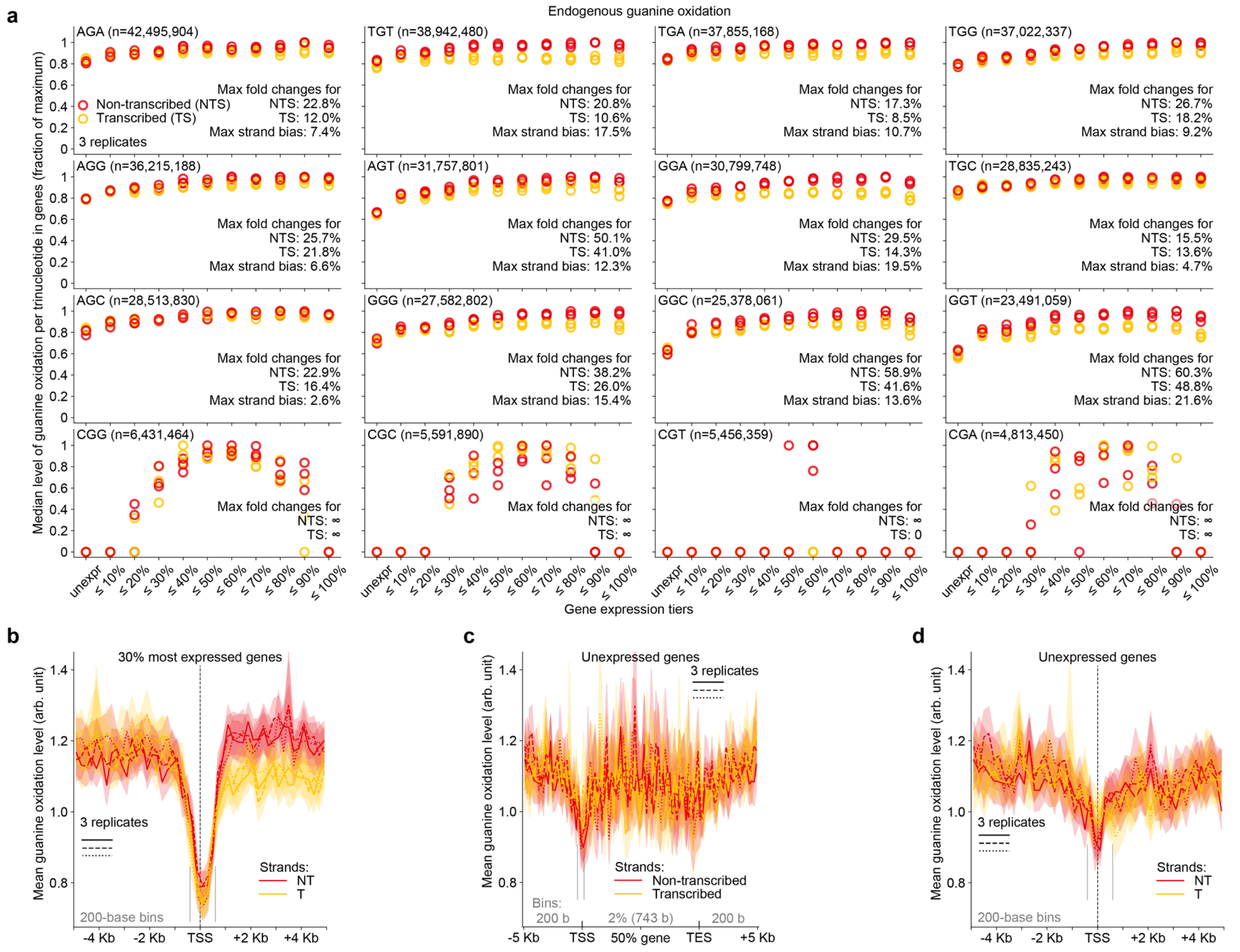


**Supplementary Fig. 4 | Oxidized guanines accumulate in gene bodies with increasing gene expression and show a strand bias.** **a** Median level of endogenous guanine oxidation in protein-coding genes considered separately for individual trinucleotide contexts and strands versus gene expression in HAP1 cells. The level of trinucleotide-specific guanine oxidation is normalized by the count of the respective trinucleotide within each protein-coding gene in the reference genome. For each replicate, the level of guanine oxidation is further normalized by the maximal value among the medians of gene-expression tiers and strands (for each sample, the maximal – among all tiers and strands – trinucleotide-specific value is set to 1; see normalization formulas in **Methods**). The trinucleotides are sorted according to the total occurrence within the protein-coding genes in the reference genome, with respective numbers indicated in parentheses. Maximal fold change for a strand: we averaged the replicate values at each tier, related these averages to the average of the unexpressed genes and indicated the maximal percentage of the change. Maximal strand bias: for every tier starting from ≤40%, we related the averages of the replicate medians between the non-transcribed and transcribed strands and indicated the maximal percentage of the change. **b-d** Strand-specific profile of the mean guanine-oxidation level and its 95% c.i. throughout the gene body and its upstream and downstream regions in highly expressed (**b**: n=3,994) and unexpressed (**c-d**: n=3,428 genes); solid, dashed, and dotted curves with shades: different biological replicates; vertical dashes demark TSS-adjacent regions analyzed in Fig.4m-p; genes are aligned only at TSS (**b,d**) or at both TSS and TES (**c**). Arb. unit: **Methods** describe guanine-oxidation level normalization. NT: non-transcribed, T: transcribed. TTS and TES: transcription start and end sites. Kb: kilobase; b: base.

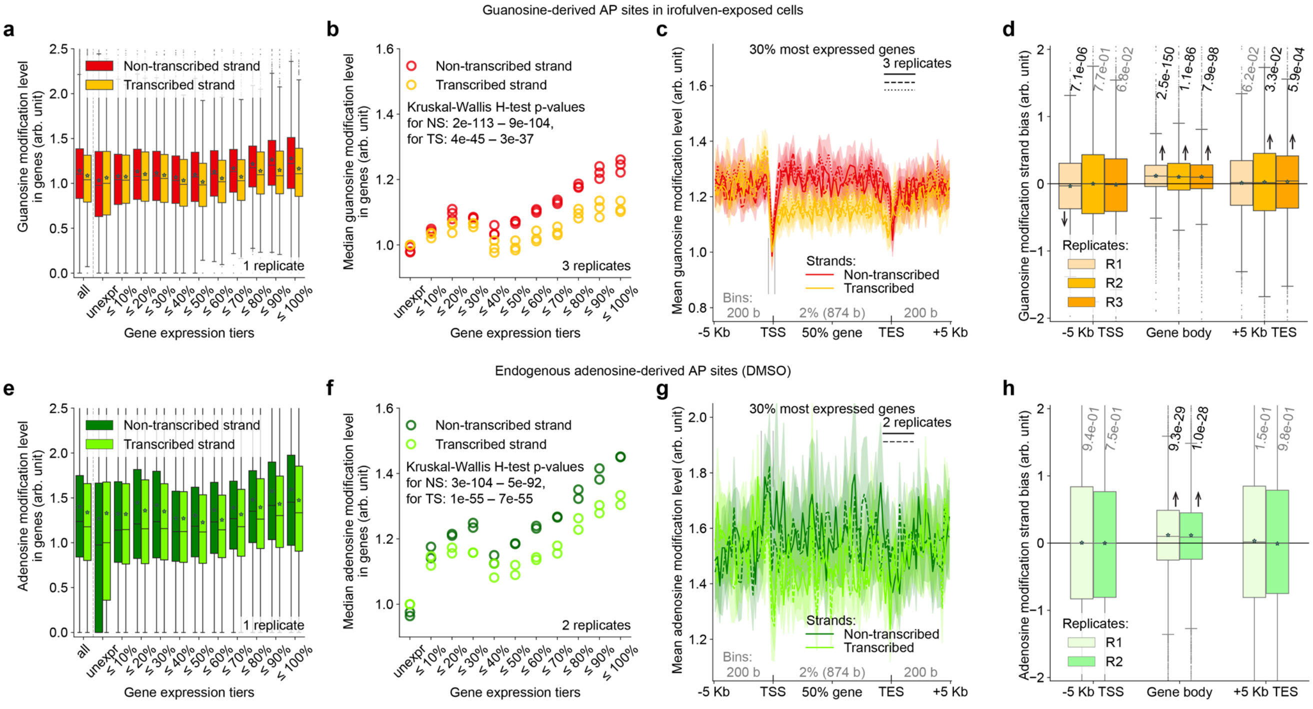


**Supplementary Fig. 5 | AP sites accumulate in gene bodies with increasing gene expression and show a strand bias. a-d** Guanosine-derived AP sites in irofulven-exposed U2OS cells throughout gene bodies and adjacent regions. **e-h** Endogenous adenosine-derived AP sites in vehicle-(DMSO)-exposed U2OS cells throughout gene bodies and adjacent regions. The plots are built analogously to Fig. 4e-l.

**a**

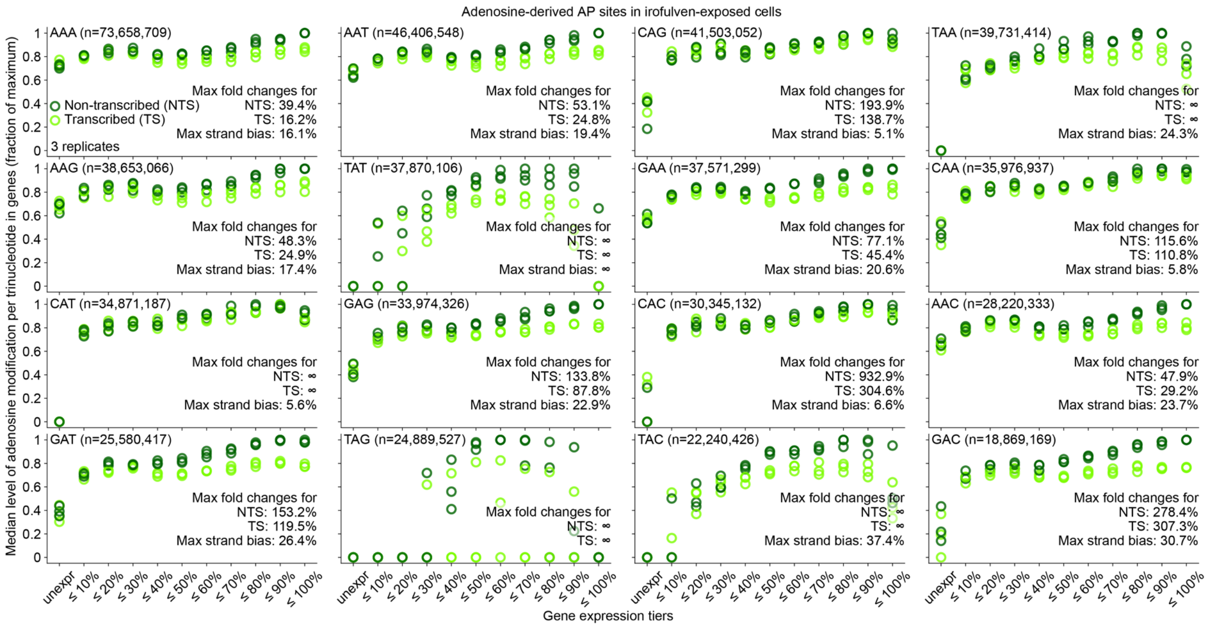

**b**

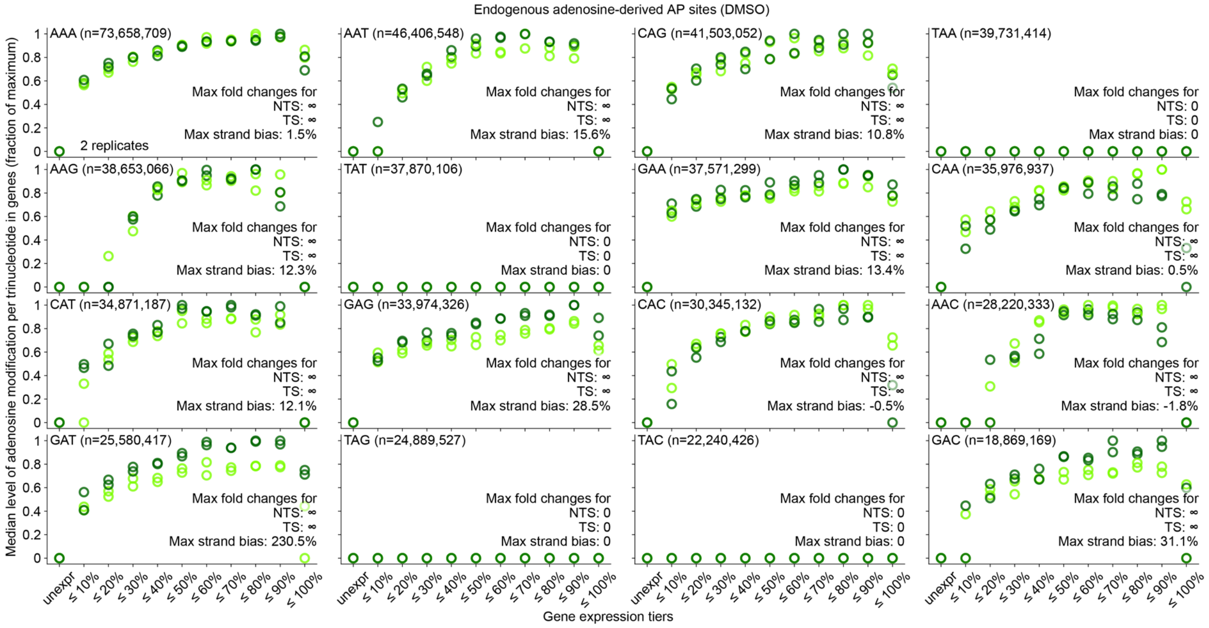

**Supplementary Fig. 6 | Adenosine-derived AP sites accumulate in gene bodies with increasing gene expression and show a strand bias. a-b** Median level of adenosine-derived AP sites in irofulven-exposed (**a**) or vehicle-(DMSO)-exposed (**b**) U2OS cells in protein-coding genes considered separately for individual trinucleotide contexts and strands versus gene expression in unexposed U2OS cells. The plot is built analogously to Supplementary Fig. 4a.

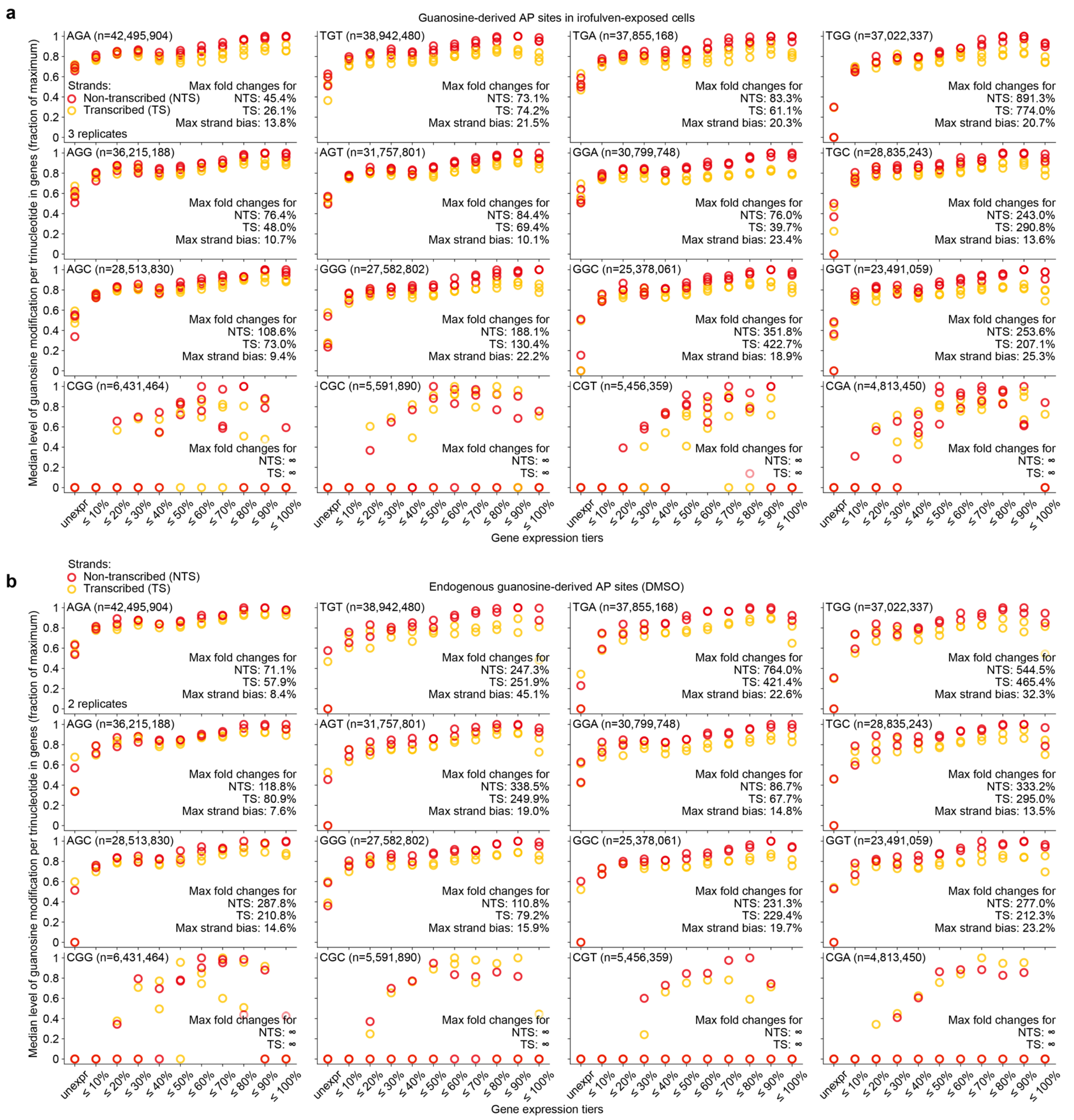


**Supplementary Fig. 7 | Guanosine-derived AP sites accumulate in gene bodies with increasing gene expression and show a strand bias. a-b** Median level of guanosine-derived AP sites in irofulven-exposed (**a**) or vehicle-(DMSO)-exposed (**b**) U2OS cells in protein-coding genes considered separately for individual trinucleotide contexts and strands versus gene expression in unexposed U2OS cells. The plot is built analogously to Supplementary Fig. 4a.

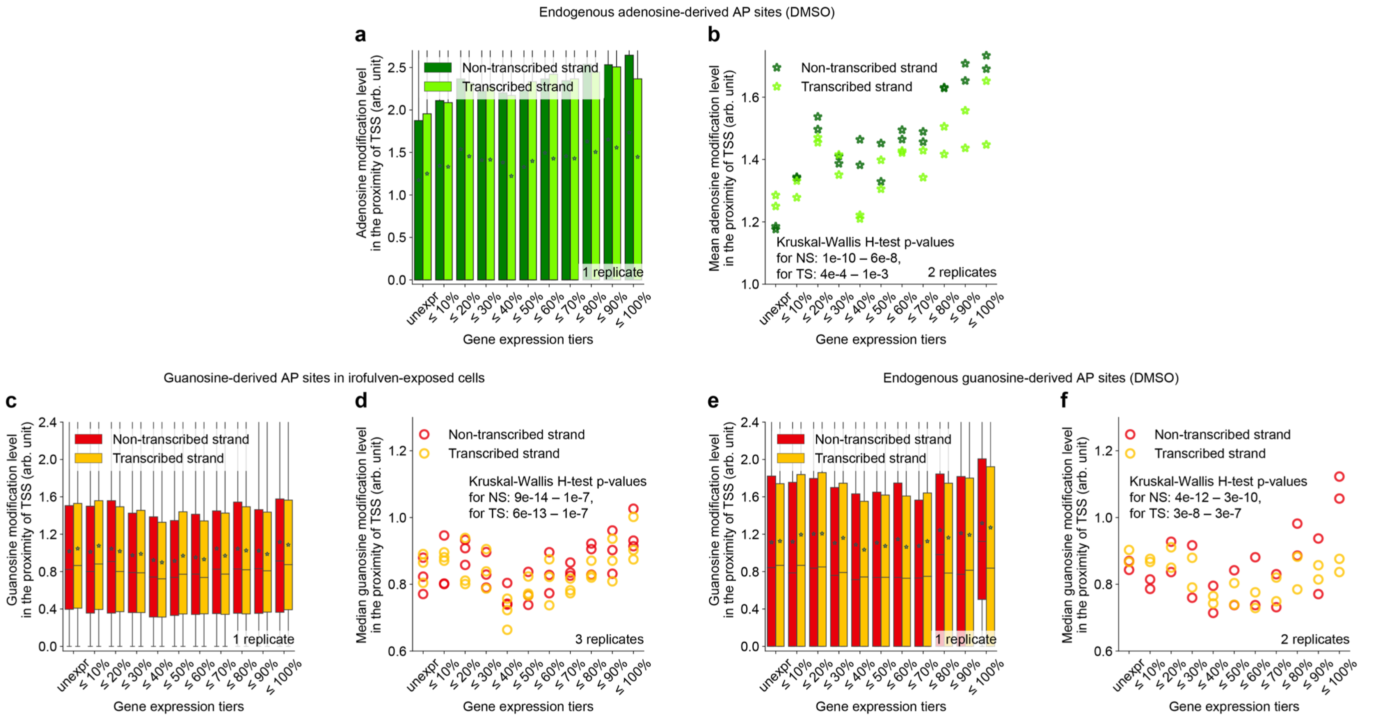


**Supplementary Fig. 8 | DNA-modification level around the TSS in each strand in protein-coding genes versus gene expression level. a-b** Adenosine-derived AP sites in vehicle-(DMSO)-exposed U2OS cells between -1000 and 0 b relative to TSS. **c-f** Guanosine-derived AP sites in irofulven-exposed (**c-d**) or vehicle-(DMSO)-exposed (**e-f**) U2OS cells between -400 and +600 b relative to TSS. The plots are built analogously to Fig. 4m-p.

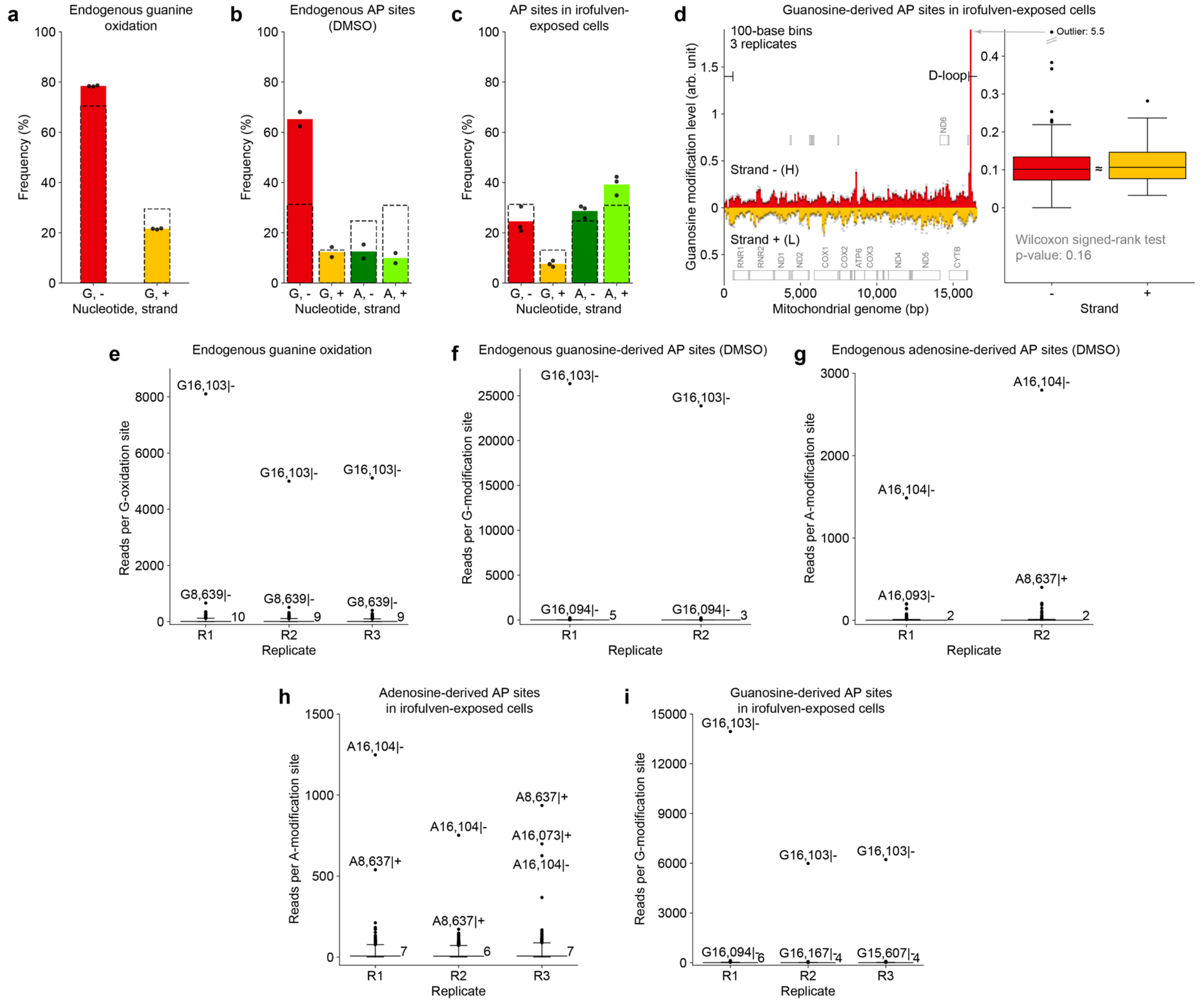


**Supplementary Fig. 9 | The distribution of DNA modifications in mtDNA. a-c** The distribution of endogenous DNA oxidation in HAP1 cells (**a**), AP sites in vehicle-(DMSO)-exposed U2OS cells (**b**) and AP sites in irofulven-exposed U2OS cells (**c**) throughout the mtDNA strands and affected nucleosides; markers, replicates; filled bars, means. Dashed bars, the distribution of respective nucleotides according to the reference genome. **d** Left panel: The strand-specific profile of guanosine depurination in irofulven-exposed U2OS cells throughout mtDNA. Right panel: bin-specific mean values from the left panel aggregated within each strand. Wilcoxon signed-rank test: 166 pairs of strand-specific values, two-sided alternative. The plots are built analogously to Fig. 5. **e-i** The distribution of the number of unique reads (with individual RICs) supporting a DNA modification site in mtDNA across the replicate experiments. Distribution plot from the bottom to the top: short horizontal line, $1^{st}$ percentile; long horizontal line, median, the value is provided; short horizontal line, $99^{th}$ percentile; markers, values bigger than the $99^{th}$ percentile. The coordinates of the sites with highest read coverages are provided.

### Supplementary Table

**Supplementary Table 1 | DNA oligonucleotides used in the work.** 5′–3′ orientation; oxoG, 8-oxoG; THF, tetrahydrofuran; Phos; phosphorylated; *, phosphothioester linkage; $N_3$; azido; TEG, tetra-ethylene-glycol linker; N, randomized position.

| Name | Sequence |
|---|---|
| IL1-T30 | ATCTTTTGTAGCATACATTGAAGATGTGGC |
| IL-1 | GCCACATCTTCAATGTATGCTACAAAAGAT |
| IL-1_oxoG | GCCACATCTTCAAT***oxoG***TATGCTACAAAAGAT |
| IL-1_THF | GCCACATCTTCAATGT***THF***TGCTACAAAAGAT |
| NEB_P7 | AGACGTGTGCTCTTCCGATCTAGAAGGCCTAG*****T |
| NEB_P5 | **Phos**-CTAGGCCTTCTAAGGAGATGTTGATGTGCTGC |
| MoDIS_1 | **$N_3$**-TNNNNNNNNNNCAACAAAGATCGGAAGAGCGTCGTGTAGG-***TEG***-Biotin |
| MoDIS_2 | **$N_3$**-TCAACAANNNNNNNNNNNAGATCGGAAGAGCGTCGTGTAGG-***TEG***-Biotin |
| Pr-P7 | AGACGTGTGCTCTTCCGATCTA |
| exUMI | CCTACACGACGCTCTTCCGATC |
| i5/i7 indexing primers | These sequences are from the NEBNext® Multiplex Oligos for Illumina® (Index Primers Set 1) (NEB E7335S) and can be found in the respective product manual. |

### Supplementary Note

**Supplementary Note 1 | AP-site strand bias in trinucleotide contexts within gene bodies.** Analyzing median AP-site levels in individual trinucleotide contexts (Supplementary Fig. 6 and 7), we observed elevating DNA-modification burdens with increasing gene expression for almost all trinucleotides for both irofulven-induced and endogenous depurination. Yet, the trinucleotides were found to vary in the maximal fold changes, spanning, in case of irofulven-induced damage, from 39.4% for in AAA up to infinity (elevation from zero) in TAN, CAT (Supplementary Fig. 6a) and CGN (Supplementary Fig. 7a) in the non-transcribed strand. Similarly, the strand biases proved to vary across the trinucleotides from no pronounced strand differences of irofulven-induced depurination in CAN (Supplementary Fig. 6a) and CGN (Supplementary Fig. 7a) along the gene-expression tiers to over 30% difference between the non-transcribed and transcribed strands for GAC and TAT in the tier of the most expressed genes (Supplementary Fig. 6a). Supporting that the endogenous adenosine-derived AP sites do not confound the mapping of irofulven-induced ones, we noticed different gene-expression-dependent patterns at individual trinucleotides in vehicle-treated samples compared to irofulven exposure (Supplementary Fig. 6b vs 6a), with fewer trinucleotides showing the strand bias and with generally higher fold changes as the background adenine depurination is mostly zero in unexpressed genes (Supplementary Fig. 6b). For example, in AAA context, there is a pronounced strand bias under irofulven exposure (Supplementary Fig. 6a), although vehicle exposure is associated with no difference between the gene strands (Supplementary Fig. 6b), which may be explained by polymerase stalling at irofulven-adenosine adduct in the first case and a lack of polymerase stalling at the AP site in the second case. On the contrary, trinucleotide-specific guanosine-derived AP-site levels were more similar between irofulven- and vehicle-exposed samples (Supplementary Fig. 7a-b), which, given lower rate of guanosine modification by irofulven (Fig. 2e), could potentially reflect a higher confounding effect of endogenous guanosine depurination for mapping this DNA modification in irofulven-exposed samples.
